# Bacterial capture and lysis on diatom spines reveal a suspension-feeding strategy

**DOI:** 10.64898/2026.09.07.749814

**Authors:** M. Forget, F. Wollweber, E. Case, R.J. Henshaw, N. Blitvic, J. Slomka, M. Pilhofer, J.H Hehemann, R. Stocker

## Abstract

Phytoplankton contribute nearly half of global primary production and play a central role in ocean biogeochemistry. Although traditionally viewed as phototrophs, a growing body of evidence indicates that many phytoplankton are mixotrophic, supplementing photosynthesis with heterotrophic acquisition of nutrients and organic matter. Yet, the mechanisms enabling heterotrophic feeding in diatoms remain poorly understood. Here, we show that the chitinous spines of the diatom *Conticribra weissflogii* can serve as attachment sites for the bacterium *Marinobacter adhaerens*. Cryo-electron tomography and viability staining revealed that a substantial fraction of spine-associated *M. adhaerens* undergoes lysis. Consistent with bacterial lysis on the spines, Raman spectroscopy showed that diatom cells bearing spine-associated bacteria acquired more bacteria-derived nitrogen than uncolonized cells. Although microfluidic assays revealed a low bacterial affinity for the spines, our results suggest that attachment can be promoted by fucoidan deposits identified on the spines — a compound for which bacteria showed significantly higher binding affinity than chitin — as well as by their geometry, which encounter rate calculations show to be particularly efficient at intercepting bacteria. Together, these findings reveal a previously undescribed form of suspension-feeding in phytoplankton, reminiscent of the role of zooplankton pseudopods in prey capture. Diatom spines intercept bacteria from the surrounding water, retain them near the cell surface, and facilitate subsequent nutrient acquisition, probably through bacterial lysis. Our results expand the functional repertoire of diatom spines beyond defense and buoyancy, identifying spine-mediated bacterial capture as a previously overlooked pathway contributing to phytoplankton mixotrophy.

**Significance statement:** Diatoms produce long chitinous spines whose functions have been linked to buoyancy control and defense against grazers. Here, we reveal a previously unrecognized function of these appendages: the spines of the diatom *Conticribra weissflogii* serve as attachment sites for the bacterium *Marinobacter adhaerens*, a substantial fraction of which undergoes lysis on the spine surface. Using ^15^N isotopic labelling and Raman spectroscopy, we demonstrate that diatoms harboring spine-associated bacteria acquire significantly more bacterial-derived nitrogen than uncolonized cells, consistent with spine-mediated bacterial capture contributing to diatom nutrient acquisition. We further show that *C. weissflogii* spines are coated with fucoidan, which likely promotes bacterial attachment. This suspension-feeding strategy — in which spines coated with fucoidan, intercept bacteria from the surrounding water, analogous to the slime-coated pseudopods of radiolarians and foraminiferans — has never previously been documented in phytoplankton. Our findings expand the known functional repertoire of diatom spines and identify a previously overlooked pathway of mixotrophic nutrient acquisition in phytoplankton with potentially broad implications for marine nutrient cycling.

## Introduction

Phytoplankton are a diverse group of photosynthetic microorganisms, including microalgae such as dinoflagellates, diatoms and coccolithophores, as well as cyanobacteria such as *Prochlorococcus* and *Synechococcus*. Through their photosynthetic activity and high turnover rate^1^ – exemplified by blooms visible from space^2^– they contribute approximately half of Earth’s net primary production^3^, forming the foundation of marine food webs and driving major marine biogeochemical cycles^3^. Sustaining this productivity requires efficient acquisition of essential nutrients whose availability and chemical forms strongly constrain phytoplankton growth, productivity, and geographical distribution^4^. Phytoplankton interact closely with associated bacteria that can provide essential metabolites including vitamin B12^5^ and growth-promoting hormones^6^. These interactions often involve reciprocal exchanges, whereby bacteria benefit from phytoplankton-derived organic carbon while supporting phytoplankton metabolism through the provision of key compounds^5,7^. Phytoplankton-bacteria interactions therefore play central roles in marine primary production, carbon export, and nutrient regeneration^8^. Increasing evidence also indicates that mixotrophy—the ability to combine phototrophy with heterotrophic organic matter acquisition—is widespread among phytoplankton^9–14^. Phytoplankton may thus potentially benefit from bacteria not only through metabolic exchanges with living cells, but also as a source of organic matter.

Phytoplankton exhibit remarkable morphological diversity, spanning cell sizes from a few micrometers to several hundred micrometers, and encompassing a wide range of cellular architectures, including siliceous frustules and calcium carbonate plates, motile and non-motile forms, and solitary or chain-forming lifestyles^15–17^. A recurrent feature across diverse phytoplankton lineages is the presence of long and rigid appendages radiating from the cell surface, hereafter referred to as spines (also termed setae or threads in some taxa). Spines are found in numerous genera of green algae^18,19^, cocolithophorous^20^, dinoflagellates^21^, and diatoms^22^. They can be either extensions of the cell wall or, as in the diatom genera *Thalassiosira* and *Cyclotella*, chitin fibrils extruded through specialized pores^23^. Phytoplankton spines are typically slender and can be several tens of micrometers long. Despite their frequent occurrence and striking morphology, the ecological functions of phytoplankton spines remain incompletely understood. They have primarily been interpreted as adaptations that reduce sinking rate by increasing the cell surface area (and thus the hydrodynamic drag) with minimal mass addition^24–26^ and/or to deter grazers by considerably increasing the effective cell size^27,28^. However, their potential role in mediating bacteria–phytoplankton interactions, a key component of phytoplankton ecology, remains unexplored. Because phytoplankton spines extend into the phycosphere —the micrometer-scale chemical microenvironment surrounding individual phytoplankton cells where metabolite exchanges with associated bacteria occur^29–31^— they could increase the probability of bacterial encounter and provide surfaces for bacterial attachment, analogous to the feeding appendages of some zooplankton^32^.

Motivated by scattered observations of bacteria attached to diatom spines, —including *Vibrio parahaemolyticus* on *Conticribra weissflogii*^33^, Bacteroidetes on *Chaetoceros*^34^, and *Phaeobacter spp.* on *Thalassiosira rotula*^35^, we asked whether and how these structures contribute to bacteria–phytoplankton interactions. To address this question, we used the centric diatom *Conticribra weissflogii (formerly Thalassiosira weissflogii)*, which produces dozens of slender chitinous spines approximately 100 nm in diameter and extending several tens of micrometers from the cell surface^23^, together with the marine γ-proteobacterium *Marinobacter adhaerens*. *M. adhaerens* is a ubiquitous, motile marine γ-proteobacterium whose interaction with *C. weissflogii* has become a model system for investigating the ecological and biophysical mechanisms governing diatom–bacteria interactions^36–40^. Using electron and fluorescence microscopy, we documented *M. adhaerens* attachment to *C. weissflogii* spines and quantified the prevalence of this interaction mechanism. We then combined cryo-electron tomography and Raman spectroscopy to show that spine-associated bacteria frequently undergo lysis and that *C. weissflogii* cells colonized by *M. adhaerens* on their spines exhibit enhanced acquisition of bacteria-derived nitrogen. Finally, we explored the mechanisms governing spine colonization by combining microfluidic assays, targeted staining and encounter-rate calculations. We found that colonization results from the combination of an overall low bacterial affinity for the spines, the presence of diatom fucoidan on the spine favoring bacterial attachment, and frequent bacteria–spine encounters. Together, our results identify diatom spines as previously unrecognized structures mediating bacterial capture and reveal a previously undescribed form of suspension-feeding that provides a pathway for mixotrophic nutrient acquisition in diatoms.

### I. The bacterium *Marinobacter adhaerens* attaches to the chitin spines of the diatom *Conticribra weissflogii*

Triplicate axenic cultures of *C. weissflogii* at a concentration of 10^4^ diatoms mL⁻¹ were inoculated with the *M. adhaerens* strain Hp15^36^ expressing a red fluorophore (Dsred) at a concentration of 5×10⁵ bacteria mL⁻¹. At days 6 and 14 post-inoculation, the diatom fraction was isolated by filtering co-cultures through an 8 µm membrane and imaged by epifluorescence microscopy (20× objective). We observed that diatom cells had bacteria located several micrometers away from their surface. Those bacteria were static. Bright-field imaging revealed that they were attached to the spines of the diatom cells (Fig. 1B) and this observation was confirmed by scanning electron microscopy (SEM; Fig. 1C, Fig. S1). Spine-attached bacteria showed no apparent preferential location along the spines, attaching from the base near the cell wall to the distal tip (Fig. 1B, Fig. S1). SEM (Fig. 1B, Fig. S1) and cryo-electron tomography imaging (cryoET; Fig. S2) further revealed that bacteria attached to diatom spines had structures consistent with pili. Although bacterial pili have been hypothesized to contribute to phytoplankton surface colonization^41^, this study provides, to our knowledge, the first microscopic observation of pili-like appendages on bacteria attached to a phytoplankton cell. In 14-day old cocultures, epifluorescence microscopy showed that bacteria were also observed attached directly to the diatom cell wall, as previously reported for this association^37^ (Fig. 1F), representing 31% ± 7.3 of all attached bacteria. This cell-wall attachment mode was not detected in exponential-phase cocultures, suggesting that either diatom surface properties or bacterial attachment behavior changes as cocultures age.

**Figure 1.**
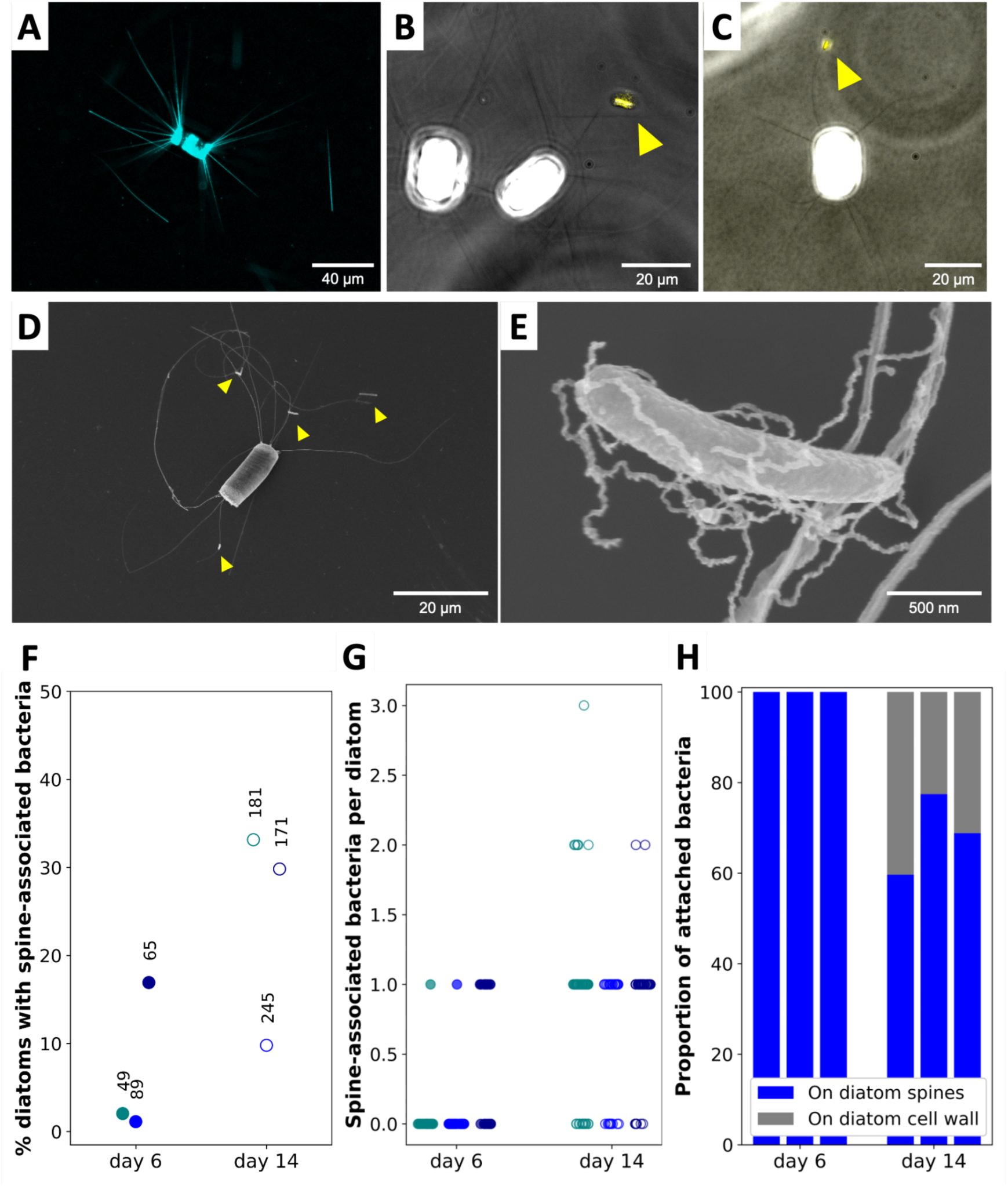
*M. adhaerens* attaches to *C. weissflogii* spines in cocultures. **A,** Calcofluor White staining of C. weissflogii spines. **B-C**, Bacteria (indicated by yellow arrowheads) attached to diatom spines. **D**, Scanning electron microscopy image of a diatom with multiple spine-associated bacteria and **E**, a magnified image showing bacterial pili entwined with a spine. **F–H**, Quantification of bacterial colonization of diatoms in co-culture, showing the percentage of diatoms with spine-associated bacteria (F), the number of spine-associated bacteria per diatom (G), and the percentage of colonized diatoms with bacteria attached to spines (blue) vs. to the cell wall (gray) (H). Data come from three independent cocultures, each assayed at days 6 and day 14 post-inoculation. Sample sizes are indicated above each datapoint in panel F.

Epifluorescence microscopy revealed that colonized diatoms harbored one to three spine-attached bacteria (Fig. 1E). The proportion of diatom cells with spine-attached bacteria was low (6.70% ± 7.24) at day 6 post coculture inoculation, but increases markedly at day 14, reaching 24.26% ± 10.31 of the diatom cells (Fig. 1D). This result indicates that a substantial fraction of the diatom population has bacteria on its spines and that this interaction mechanism could thus represent a functionally relevant interaction from the diatom perspective.

### II. *M. adhaerens* attachment to *C. weissflogii* spines is associated with bacterial damage

CryoET analysis of *M. adhaerens* attached to *C. weissflogii* spines showed pronounced signs of bacterial damage (Fig. 2A–C). These include the release of membrane vesicles (Fig. 2A-C), a feature commonly associated with bacterial stress response^42^, and bacterial lysis. Bacterial lysis was evident for several cells from the loss of their outer membrane and overall structural integrity, their enlarged periplasmic spaces, and the release of large membrane vesicles (Fig. 2B-C). In one notable example, a spine tip physically appeared to penetrate a bacterium undergoing lysis (Fig. 2C). None of these stress-associated features were observed in the 8 tomograms of *M. adhaerens* cells from a diatom-free culture, suggesting that they were specifically associated with the diatom–bacterium interaction. We next quantified the proportion of damaged bacteria among *M. adhaerens* cells attached to *C. weissflogii* spines in cocultures. Triplicate 14-day cocultures were filtered through an 8 µm membrane to isolate the diatom-containing fraction (>8 µm), and damaged bacteria within this fraction were identified by Sytox Green staining, a nucleic acid dye that selectively labels cells with compromised membranes (Fig. 2D-E). Across the three independent cocultures, we analyzed 51, 65, and 23 spine-associated bacteria by microscopy. Of these, 33%, 20%, and 26% (respectively) stained positive for Sytox Green, indicating that approximately one-fifth to one-third of bacteria attached to diatom spines displayed membrane damage consistent with lysis.

**Figure 2.**
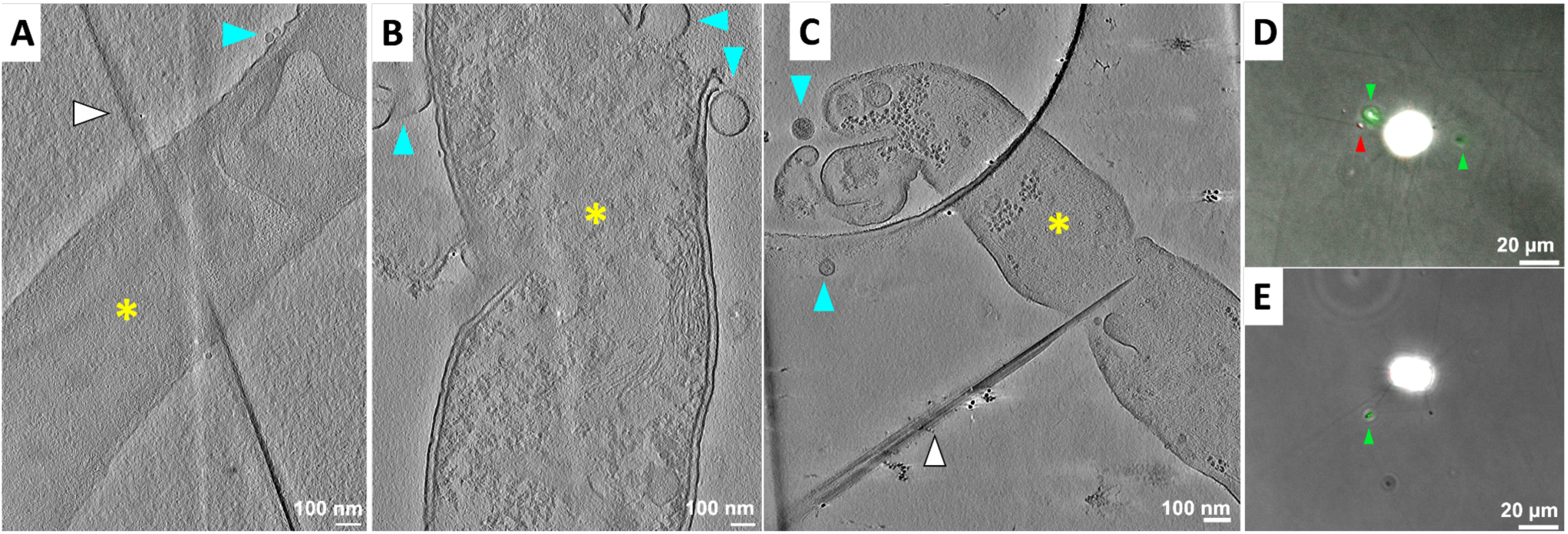
Attachment of bacteria to diatom spines is associated with bacterial damage. **A–C:** Cryo-tomographic slices of *M. adhaerens* cells (yellow asterisc) attached to *C. weissflogii* spines (white arrowheads). In panel B, the spine is not visible in the selected z-plane because it is located at a different depth within the reconstruction, above the bacterium. Attached bacteria frequently release vesicles (cyan arrowheads) and show advanced lysis with loss of cell integrity (B, C). A putative direct puncture by the spine tip was observed in one lysis event (C). **D, E:** Epifluorescence image of DsRed-tagged *M. adhaerens* on *C. weissflogii* spines after staining with the viability dye Sytox Green, overlaid with a bright field image of the diatom. Red, chlorophyll autofluorescence and DsRed-labeled *M. adhaerens* cells; green, Sytox-positive bacteria, indicating loss of membrane integrity. Green arrowheads: Sytox-positive bacteria, red arrowheads: Sytox-negative bacteria.

Lysis and vesicle release by bacteria attached to diatom spines are expected to increase the availability of bacteria-derived nutrients in the immediate vicinity of the diatom cell wall. We therefore investigated whether diatoms harboring spine-associated bacteria exhibit enhanced uptake of bacteria-derived nutrients compared to uncolonized diatom cells using Raman spectroscopy.

### III. *C. weissflogii* cells harboring attached bacteria on their spines uptake more bacteria-derived nitrogen

Diatoms were cultured for 6 days under three conditions: (1) axenically in standard f/2 medium containing ^14^NO₃⁻ (negative control, “14N-diatoms”), (2) axenically in f/2 medium containing ^15^NO₃⁻ (positive control, “15N-diatoms”), and (3) in coculture with ^15^N-labeled *M. adhaerens* in f/2 medium lacking NO₃⁻.

First, to determine how ^15^N incorporation affects the Raman signature of *C. weissflogii*, we compared spectra of ^14^N-diatoms and ^15^N-diatoms. Across three independent cultures per treatment, with Raman spectra collected from 25 diatom cells per culture, ^15^N incorporation induced significant downshifts at four Raman peaks that correspond to nitrogen-containing moieties in chlorophyll^43–45^(Fig. S3). Among these, we selected the peak centered at 916 cm⁻¹ in ^14^N-diatoms which shifted to 913.8 cm⁻¹ in ^15^N-diatoms (two-sample t-test, p = 0.015, Fig. 3A), because it exhibited the largest shift and thus provided the most sensitive indicator of ^15^N incorporation.

**Figure 3.**
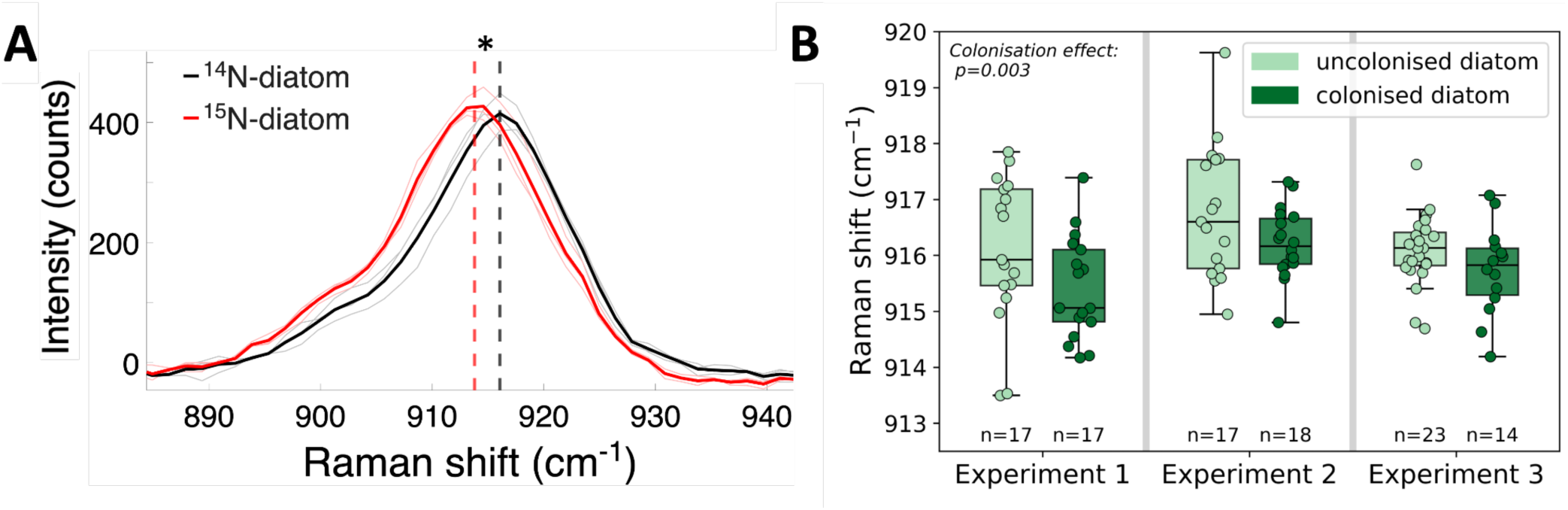
Raman spectroscopy reveals enhanced uptake of bacterial nitrogen in colonized diatom cells relative to uncolonized cells. **A:** Raman spectra of diatoms grown in ^14^N f/2 medium (black) and ^15^N f/2 medium (red), showing the downshift of the 916 cm⁻¹ Raman peak along with ^15^N incorporation. The peak corresponds to a chlorophyll-associated macrocycle vibration involving C-C-N skeletal bending. Thin transparent lines represent the mean Raman spectra obtained from 25 individual diatom cells in each of three independent cultures per treatment. Thick lines represent the average spectrum across the three biological replicates for each treatment. Vertical dashed lines indicate the mean position of the 916 cm⁻¹ peak for each treatment. Statistical comparison (t-test): * p-value < 0.05. **B:** Raman shift of the 916 cm^-1^ peak in uncolonized diatoms (light green) and in diatoms with *M. adhaeren*s bacteria attached to their spines (dark green) across three biological replicates. n indicates the number of diatoms for which the Raman signal was measured.

We then used the position of this peak as a relative indicator of ^15^N enrichment to compare nitrogen incorporation among individual *C. weissflogii* cells. We applied this approach to diatoms from the coculture treatment to determine whether spine colonization influenced *M. adhaerens*-derived nitrogen acquisition. Prior to Raman measurements, individual *C. weissflogii* cells were examined by microscopy (60× objective) and classified according to their colonization status as either colonized (typically bearing 1–2 spine-attached *M. adhaerens* cells) or uncolonized. Diatoms with cell-wall-attached bacteria, which were rarely observed in these cocultures, were excluded from the analysis. Raman spectra were subsequently collected from individual cells and processed for smoothing and baseline subtraction (see Materials and Methods). Finally, the position of the 916 cm⁻¹ peak was quantified for each *C. weissflogii* cell. Across three independent coculture experiments, colonized diatoms consistently exhibited a downshift of the 916 cm⁻¹ Raman peak relative to uncolonized diatoms from the same coculture (Fig. 3B), indicating greater relative ^15^N incorporation in cells bearing spine-attached bacteria. To statistically evaluate this effect while accounting for baseline differences among experiments, we fitted a linear model with colonization status and experiment as categorical predictors. Because Raman-shift variability differed among colonization status, statistical inference was performed using HC3 heteroskedasticity-consistent standard errors^46^. This analysis revealed a significantly lower 916 cm⁻¹ peak position in colonized compared with uncolonized diatoms after accounting for experimental variation (one-sided HC3-based test, p = 0.003). An independent experiment-level analysis yielded the same conclusion (one-sided t-test, p = 0.016; see Materials and Methods). Overall, these results demonstrate that bacterial colonization of diatom spines is associated with enhanced uptake of bacteria-derived nitrogen by diatom cells. Given that a substantial proportion of *C. weissflogii* cells exhibit spine-associated bacteria in 14-day-old cocultures (Fig. 1D), this interaction may represent a widespread and ecologically relevant pathway for bacteria-derived nutrient acquisition within diatom populations.

We next investigated the physical and chemical determinants of *M. adhaerens* attachment to *C. weissflogii* spine.

### IV. Low bacterial affinity for diatom spines and potential fucoidan-associated attachment sites

Under coculture conditions, we estimate that the spines of a diatom cell encounter an average of 400 bacteria per hour at day 6 post inoculation (Note S3). This high encounter rate contrasts with the low number of *M. adhaerens* cells observed on *C. weissflogii* spines under these conditions (Fig. 1F-G) which suggests that bacterial attachment is constrained by biochemical compatibility between bacterial surfaces and diatom spines. We developed a microfluidic setup to quantify bacterial affinity for attachment to the spines of *C. weissflogii* to test this hypothesis.

Initially axenic diatoms trapped between micropillars in a microfluidic channel (Fig. S8) were exposed to a controlled flow of a bacterial suspension (Fig. 4A,). The flow speed was set to 20 µm s^-1^ to match the swimming speed of *M. adhaerens*^47^, such that bacteria–spine encounters occurred at relative velocities approximately comparable to those expected in the ocean. Diatoms were harvested from 6-day cultures, a condition under which we found bacterial attachment to occur exclusively on the spines, rather than also on the diatom cell body (Fig. 1E). After 30 min of exposure to the bacterial suspension, the fraction of diatoms bearing spine-attached bacteria was quantified by epifluorescence microscopy. Using this quantity and the estimated number of flow-driven encounters between bacteria and diatom spines that occurred during the experiment, we computed the probability *q* that an encounter between a bacterium and a diatom spine results in bacterial attachment to the spine (detailed calculations are provided in Note S3). We considered *q* as a proxy for the bacterial affinity for attachment to the spines. We measured low bacterial affinity for *C. weissflogii* spines, with *q =* 3.7 x 10⁻⁴ for *M. adhaerens*, indicating that only approximately 1 in 2,700 bacteria–spine encounters results in attachment (Fig. 4B). Experiments using *Alteromonas macleodii* and several Vibrio species yielded values of *q* of the same order of magnitude (Fig. 4B), as did experiments using *M. adhaerens* with diatoms harvested 14 days post-inoculation (Fig. S5). The low bacterial affinity for *C. weissflogii* spines we measured is further supported by calculations of *q* derived from coculture experiments (*q =* 7.0 x 10⁻^6^, Note S3). Together, our results indicate that the low affinity of bacteria to diatom spines is a conserved feature across the bacterial species and experimental conditions examined. To understand how spine colonization nevertheless occurs despite this low overall affinity (Fig. 1, Fig. 4), we investigated the chemical composition of the spine surface to determine whether localized attachment sites with higher bacterial affinity are present.

**Figure 4.**
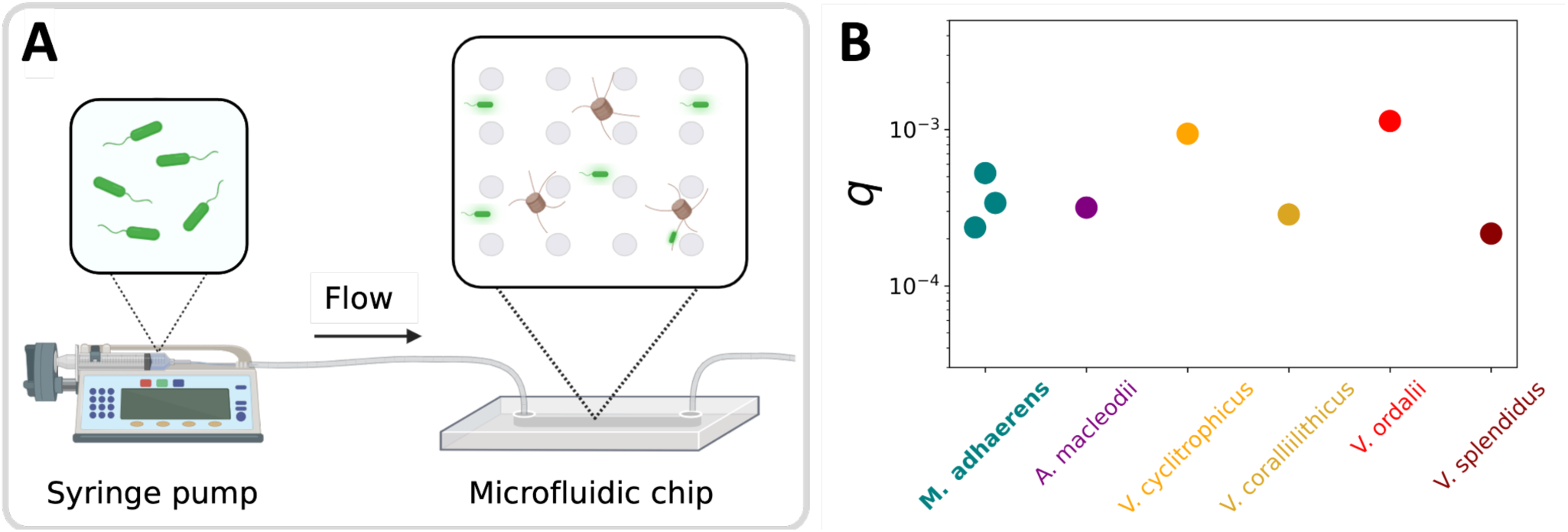
Microfluidic quantification demonstrates low bacterial affinity for attachment to *C. weissflogii* spines. **A**: Schematic of the microfluidic setup. **B**: The probability *q* that a bacterium attaches to diatom spines upon encounter, calculated from the fraction of diatoms colonized after 30 min of exposure of initially axenic *C. weissflogii* cells to suspensions of *M. adhaerens*, *A. macleodii*, or four Vibrio strains. Symbols show the results from individual experiments.

First, to probe the relative contributions of hydrophobic and electrostatic interactions to spine attachment, we performed microfluidic experiments (same setup as in Fig. 4A) using 1-µm-diameter fluorescent polystyrene beads (5×10⁶ beads mL⁻¹) as a simplified model system. Unmodified polystyrene (P) beads — which are hydrophobic — were compared with negatively charged, hydrophilic carboxylated (C). C-beads provide a proxy for bacterial cells, as they are comparable in size and have a negatively charged surface, analogous to ionized phosphate and carboxyl groups on bacterial cell walls^48^.

We quantified bead attachment by measuring the average number of spine-attached beads per diatom after 30 min of exposure to the bead suspension (Fig. 5A) and by tracking the number of spine-associated beads per diatom throughout the duration of the experiment (*i.e.*, 30min, Fig. 5B). The vast majority of attached beads were located on the spines (Fig. S6). Attachment of both bead types was 3.6-fold higher for diatoms from 14-day cultures than for those from 6-day cultures (Fig. 5A; *p* = 0.017). Because the number of spines per diatom did not vary significantly with culture age (Fig. S4), the greater number of spine-attached beads per diatom indicates that spines of diatoms from 14-day cultures have an increased propensity to bind beads. We did not detect a corresponding increase in bacterial affinity for the spines of 14-day diatoms in the microfluidic assay with live bacteria (Fig. S5). This difference may partly reflect the greater number of bead-attachment events per diatom (relative to bacterial attachment, for which we observed approximately one bacterium per diatom in the microfluidic assay), providing greater statistical power to resolve differences in affinity, as well as the fact that bacterial attachment, unlike bead binding, depends on bacterial physiology, which may vary with the physiological state of the diatom host. P-beads, which are hydrophobic, exhibited strong affinity for *C. weissflogii* spines, reaching up to 15 beads per diatom after 30 min (Fig. 5B). This observation is consistent with the hydrophobic nature of chitin and may help explain the limited colonization of diatom spines by bacteria, whose cell surfaces are generally hydrophilic^48^. C-beads, which are hydrophilic, showed a distinct accumulation dynamic relative to P-beads. A segmented regression analysis (see Material and Methods and Fig. S7) identified a breakpoint after 9 minutes of exposure to C-beads, when diatoms carry on average 4 beads per cell (Fig. 5B, Fig. S7). After this breakpoint, the slope of the fitted linear regression decreased by ∼75% and approached zero (0.04 beads min^-1^; Fig. 5B), indicating saturation of C-beads attachment. No comparable transition was observed for P-beads, whose accumulation rate remained nearly constant throughout the experiment (Fig. 5B).

**Figure 5.**
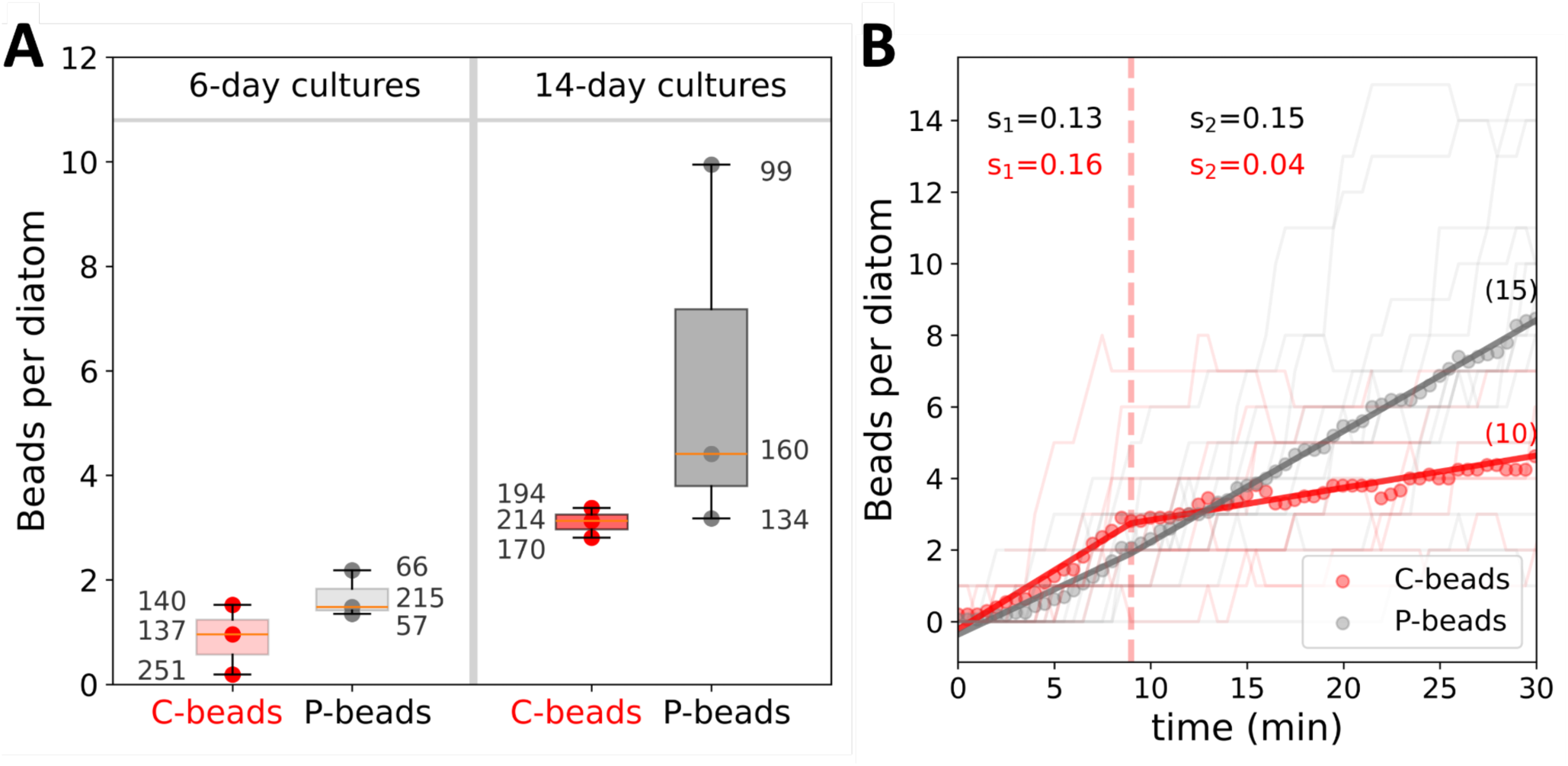
Microfluidic assays using 1-µm beads reveal limited hydrophilic binding sites on *C. weissflogii spines.* **A:** Mean number of carboxylated (hydrophilic; “C-beads”) and unmodified polystyrene beads (hydrophobic; “P-beads”) attached to the spines of single diatom cells after 30 min exposure measured in a microfluidic setup (Fig. 4A). Diatom cells came from 6-day (left) or 14-day (right) axenic cultures. Circles show the results from three independent experiments. Values next to the data points indicate the number of diatoms examined in each experiment. In each boxplot, the central line indicates the median, the box bounds the interquartile range (25th–75th percentiles) and the whiskers extend to the full data range. **B:** Time course of the number of beads attached to single diatoms over 30 min (thin lines), along with the average over 10 (P-beads) and 15 (C-beads) diatoms, respectively. Diatom cells came from a 21-day-old axenic culture. The dotted vertical red line indicates the breakpoint in C-bead colonization dynamics identified by segmented regression (see Materials and Methods). No breakpoint was identified for P-beads. Solid lines represent the linear regressions fitted before and after the breakpoint. The slope values (s_1_ and s_2_) of each fitted regression are reported next to the corresponding lines.

Together, these results suggest that only a limited number of sites on the spines support attachment of bacteria-like, carboxylated beads. These attachment sites likely become saturated after approximately four C-beads are attached per diatom, preventing further bead accumulation. SEM and cryoET analyses from this study (Fig. 1D-E, Fig. 2A and C, Fig. S1-2), together with those reported for other Thalassiosirales^35,49–51^, did not reveal any morphological features along the spines that could indicate preferential attachment sites for bacteria. We therefore investigated whether such sites could instead arise from localized chemical features of the spine surface, focusing on the potential role of diatom polysaccharides in creating hydrophilic microenvironments that promote bacterial attachment on the otherwise hydrophobic chitinous surface.

To investigate the presence of diatom polysaccharides on the spines, we employed an immunolabeling protocol^52^ with a panel of primary antibodies targeting polysaccharides previously identified on diatom surfaces, including fucoidan, (1→4)-β-D-galactoglucomannan, (1→4)-β-D-xylan/arabinoxylan and arabinogalactan protein glycan^52–54^ (Table S1). Among the polysaccharides tested, only fucoidan —a sulfated, fucose-rich and highly branched polysaccharide— produced a strong signal on *C. weissflogii* cells from a 6-day axenic culture. The signal was localized around the pores from which spines originate, forming a dotted circular pattern (Fig. 6A, inset). While fucoidan-specific antibody labeling was largely absent from the spines themselves, with only occasional signal detected at their base (Fig. 6A, inset), the strong enrichment of fucoidan around the spine extrusion pores suggests that fucoidan may be deposited onto spines during their formation. However, any fucoidan coating along the mature spines may remain below the detection limit of immunolabeling due to their extreme slenderness. We then stained diatoms from a 6-day axenic culture using Aleuria Aurantia Lectin (AAL), a lectin specific to fucose linked either (α-1,6) to N-acetylglucosamine-related or (α-1,3) to N-acetyllactosamine-related structures. Lectin staining offers improved signal amplification relative to immunolabelling. AAL staining revealed the presence of fucose-containing polysaccharides on the chitinous spines of *C. weissflogii* (Fig. 6B-E). This observation is consistent with an abundance of fucoidan on the pores through which spines are extruded (Fig. 6A), as well as with the previously reported presence of fucose-containing polysaccharides on the chitinous spines of *Thalassiosira rotula*, a close relative of *C. weissflogii*^55^. To quantify the binding affinity of *M. adhaerens* for fucoidan and further test whether fucoidan could provide hydrophilic attachment sites on diatom spines, we performed a bacterial adhesion assay. Bacteria were incubated overnight in wells of a 96-well plate coated with either chitin or fucoidan, with uncoated wells serving as negative controls. After overnight incubation, the bacterial suspension was removed, non-adherent cells were rinsed away, and bacterial attachment was quantified by epifluorescence microscopy (see Materials and Methods). *M. adhaerens* showed no preferential attachment to chitin-coated surfaces relative to uncoated controls (Fig. 7F; test *p* = 0.53). In contrast, bacterial attachment was 3.4-fold higher on fucoidan-coated surfaces (Fig. 7G; *p* = 0.0001). Together, these results indicate that although the chitinous spine surface exhibits low overall affinity for *M. adhaerens*, localized fucoidan deposits could provide favorable attachment sites for bacterial colonization.

**Figure 6.**
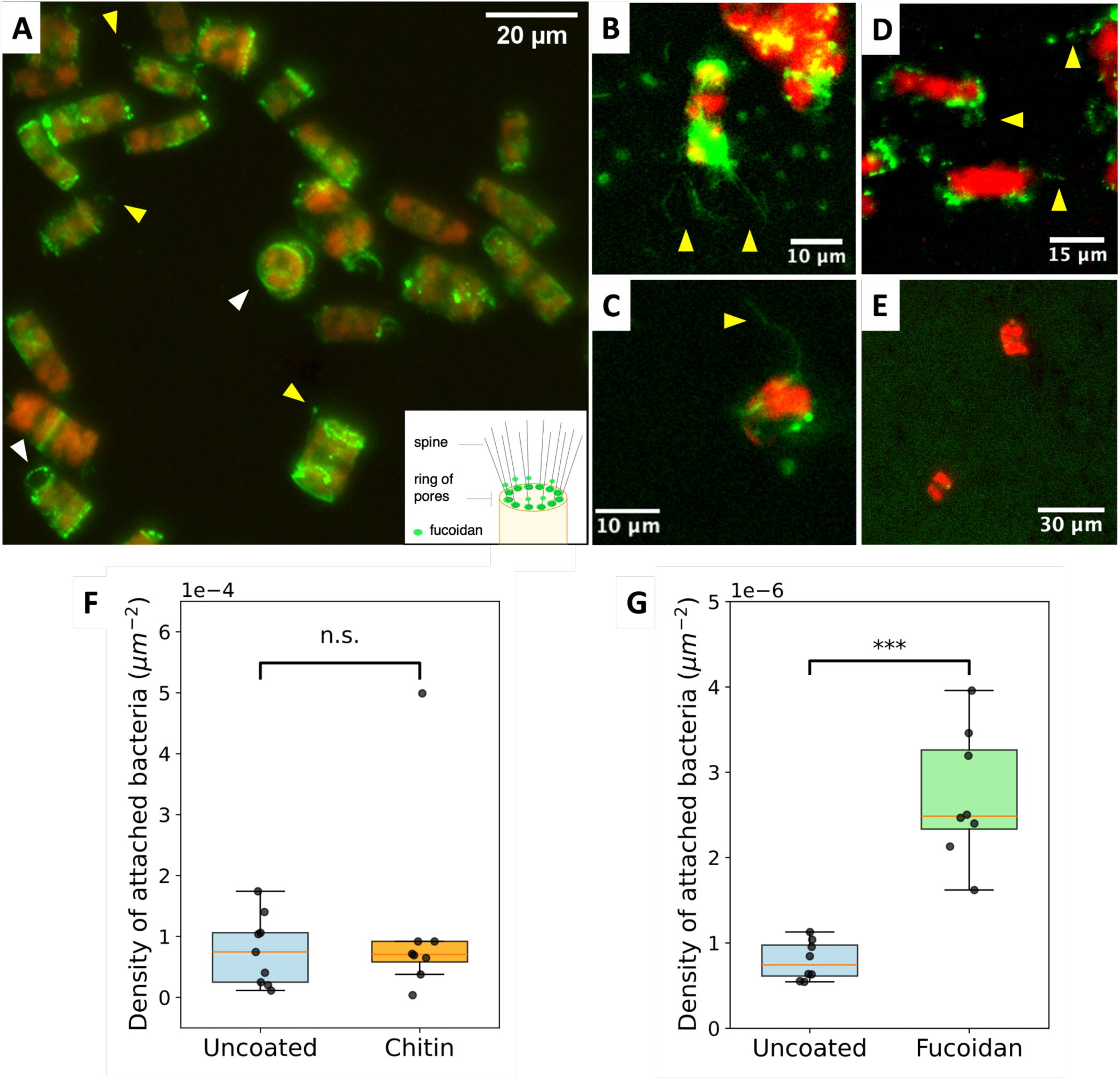
Fucoidan coating on spines mediates *M. adhaerens* attachment. **A**: Immunolabeling with a fucoidan-specific primary antibody (BAM-2) highlights a fucoidan coating at the ring of pores from which spines are extruded (white arrowheads). Yellow arrowheads indicate fucoidan signal detected at positions consistent with the location of spine bases. Green, secondary antibody conjugated to FITC bound to the fucoidan-specific primary antibody; red, diatom chlorophyll autofluorescence. The inset schematic indicates the spatial position of the fucoidan signal relative to the diatom cell and spines. **B–C**: Staining with Aleuria Aurantia Lectin, specific for fucose residues, confirms the presence of fucose-rich compounds on *C. weissflogii* spines (yellow arrowheads). Green, AAL lectin staining indicating the localization of fucose-containing polysaccharides on the diatom surface. Red, diatom chlorophyll autofluorescence. **E:** Negative control in which *Aleuria aurantia* Lectin was omitted, confirming that the green fluorescence signal observed in stained samples does not arise from autofluorescence. **F–G**: Surface density of M. adhaerens cells (cells µm^-2^) attached to the bottom surface of individual wells in a 96-well plate coated with chitin or fucoidan after overnight incubation and removal of non-attached cells (F–G). Each datapoint represents one individual well. In each boxplot, the central line indicates the median, the box bounds the interquartile range (25th–75th percentiles) and the whiskers extend to the full data range. Statistical comparisons (t-test for the experiment with fucoidan, Welch’s test for the experiment with chitin): *** p-value < 0.0005, ns non-significant.

**Figure 7.**
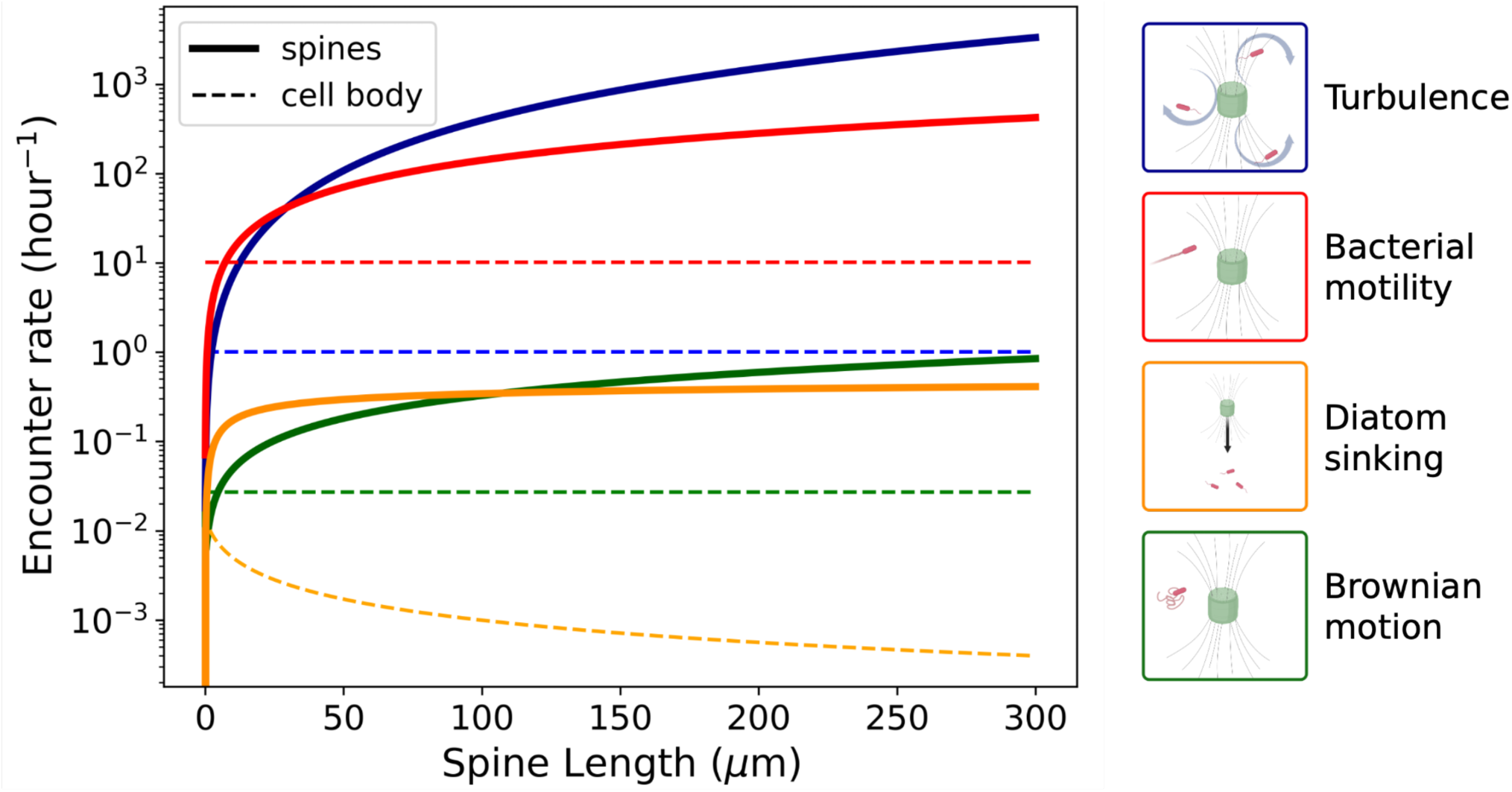
Estimated encounter rate of bacteria with a diatom that has 10 spines, for different encounter mechanisms including turbulence, bacterial motility, diatom sinking and Brownian motion (color-coding and legend on the right), as a function the length *L* of the spines. Solid lines: encounters between bacteria and the spines of the diatom. Dashed lines: encounters between bacteria and the diatom cell body. Calculations assume a turbulent kinetic energy dissipation rate of *ε* = 10⁻⁶ W kg^-1^, a bacterial swimming speed of *U* = 50 μm s^-1^, a diatom sinking velocity of *V₀* = 1 μm s^-1^, a bacterial diffusivity of *D_bact_* = 10^-13^ m² s^-1^, and a bacterial concentration of 10⁶ cells mL^-1^. Details of the calculations are provided in Note S2.

### V. Frequent bacteria–spine encounters support spine colonization

In addition to fucoidan deposits providing more favorable attachment sites than the underlying chitinous surface, bacterial colonization of *C. weissflogii* spines may be further facilitated by the high frequency of bacteria–spine encounters.

We computed encounter rates between bacteria and either the spines or the cell body of a diatom for four encounter mechanisms: turbulence, bacterial motility, diatom sinking and Brownian motion (Fig. 7, Note S2). We modeled a diatom cell with 10 spines, based on experimental measurements of spine number per cell obtained by microscopy after staining chitin with Calcofluor White (Fig. S4).

For all four encounter mechanisms, bacteria encountered the spines of the diatom cell substantially more frequently than the diatom cell body. Spines are particularly efficient at encountering bacteria in turbulence, as they extend away from the cell body into regions of higher relative flow velocity, as well as when the diatoms cells are sinking, because spines generate minimal hydrodynamic disturbance to the surrounding flow compared with the larger cell body bacteria. For 100-µm-long spines, turbulence-and sinking-driven encounter rates were approximately 400- and 350-fold higher, respectively, than those for the diatom cell body (Fig. 7).

Multiplying the probability of bacterial attachment to the spines upon an encounter (*q* = 3.7×10^4^, Fig. 4B) and assuming that turbulence, bacterial motility, sinking, and Brownian motion contribute independently to encounter rate^56^ (Fig. 7), we calculated the attachment rate of bacteria to *C. weissflogii* spines under typical bloom conditions (assuming 10⁶ bacteria mL⁻¹ and 10⁵ diatoms mL⁻¹, ref. ^57–60^, Note S4). Spine colonization is predicted to be most pronounced in the upper mixed layer where bacteria–diatom encounters are primarily driven by turbulence (Note S4). Under high turbulence (*ε* = 10⁻^5^-10⁻^4^ W kg⁻¹ ref.^61,62^), the attachment rate reaches approximately 36 bacteria per diatom per day. As diatoms sink into deeper, less turbulent waters, bacterial motility becomes the dominant encounter mechanism (Note S4). Although spines continue to intercept bacteria much more efficiently than the cell body under this encounter mechanism (Fig. 7), the attachment rate decreases by one order of magnitude to 2.4 bacteria per diatom per day (Note S4). Together, these calculations show that, despite an overall low bacterial attachment affinity, frequent bacteria–spine encounters can sustain substantial levels of spine colonization under conditions representative of natural diatom blooms.

## Discussion

### A newly identified suspension-feeding pathway of nutrient acquisition in diatoms

The capacity of *C. weissflogii* to exploit spine-attached *M. adhaerens* as a nitrogen source is strikingly reminiscent of suspension feeding by direct interception, documented across diverse zooplankton. Radiolarians and foraminiferans, for instance, deploy extracellular appendages (pseudopods) to intercept and immobilize planktonic prey^9,32,63^. In these organisms, extracellular matrix components coating pseudopods increase local adhesivity and facilitate efficient prey capture^64^. Our findings suggest that *C. weissflogii* employs a convergent strategy: fucoidan-coated spines extend into the phycosphere, intercepting *M. adhaerens* cells that would otherwise escape capture by the cell body alone. Similarly to pseudopods^65^, the geometry and hydrodynamics of diatom spines is particularly well suited to maximizing encounter rates with bacteria (Notes S2–S4), an effect amplified under turbulent conditions where spines extend beyond the cell boundary layer into regions of higher relative flow velocity. Thus, even weak bacterial–spine binding can translate into substantial bacterial capture: under bloom conditions, frequent spine–bacterium encounters are predicted to result in an average of 36 attached bacteria per diatom per day, providing a potentially meaningful flux of bacterial-derived organic matter.

The fucoidan coating of *C. weissflogii* spines is likely central to bacterial attachment, increasing local adhesivity on an otherwise hydrophobic, chitinous substrate — analogous to the extracellular matrix of foraminiferan pseudopods^64^. The small number of bacteria observed on *C. weissflogii* spines and the limited retention of carboxylated beads under flow suggest that attachment sites on the spines are spatially restricted. Discrete localization of fucoidan along the spines was, however, not visible from lectin labelling except for a small number of cells showing signal concentrated near spine bases. Further elucidating the mechanisms governing fucoidan deposition and spatial patterning on *C. weissflogii* spines represents an important avenue for understanding the regulation of *M. adhaerens* attachment. One candidate mechanism for the regulation of fucoidan distribution involves chitosan, the partially deacetylated form of chitin. Chitosan has previously been detected in the cell wall of *C. weissflogii*^66^, and the diatom genome encodes multiple chitin deacetylases^67^, suggesting that localized chitin deacetylation could generate protonated amine groups capable of forming ionic interactions with the negatively charged sulphate groups of fucoidan^68^— thereby creating attachment sites along the spine surface. Testing this hypothesis will require chitosan-specific lectin labelling that does not cross-react with chitin and would provide an important mechanistic link between chitin modification and the regulation of bacterial colonization.

Unlike suspension-feeder zooplankton, which immobilize prey on pseudopods and subsequently ingest them through pseudopodal withdrawal and phagocytosis^9,63^, diatom spines are non-retractable, necessitating alternative mechanisms of prey nutrient extraction. We observed that 26.3% (± 6.5%) of spine-associated bacteria show signs of physical damage — a proportion that likely arises from diatom–bacterium interaction rather than from preferential attachment of already-damaged cells (Note S1). Bacterial lysis at the spine surface is expected to release intracellular contents directly into the phycosphere, making bacterial-derived nutrients locally available for uptake by the diatom cell. We show that interactions with *C. weissflogii* spines and spine-associated fucoidan impose a measurable physiological cost on *M. adhaerens*, manifested as bacteriostatic effects that likely constrain bacterial proliferation on the spine surface (Note S1). Testing whether these effects contribute to lysis of spine-attached bacteria — and through what mechanisms — will require approaches capable of resolving the physiological state of individual bacteria before and after their attachment to diatom spines.

Collectively, our findings reveal a previously unrecognized function of diatom spines as suspension-feeding apparatuses — beyond their established roles in predation defense and reducing gravitational sinking — and identify a previously overlooked route of nutrient acquisition in diatoms.

### Spine-mediated suspension feeding supports diatom mixotrophy

The traditional dichotomy between photoautotrophic phytoplankton and heterotrophic zooplankton has been substantially revised over the past decade. A growing body of evidence has demonstrated that autotrophy and heterotrophy are not mutually exclusive in planktonic ecosystems, and that mixotrophy is widespread among plankton^9–12^, including diatoms^13,14^. Mixotrophy is now recognized as a fundamental feature of these ecosystems carrying important implications for models of nutrient cycling and carbon flux^10,11^. Yet the mechanisms through which phytoplankton acquire nutrients from particulate organic matter remain poorly understood. In particular, despite the ubiquity of diatom–bacterium interactions in marine environments^7^, the exploitation of bacterial biomass as a source of organic matter has received little mechanistic attention in diatoms — which have long been modelled as obligate photoautotrophs. The present study addresses this gap by providing the first direct mechanistic evidence of bacterivory in diatoms, revealing how spine-mediated bacterial capture contributes to the mixotrophic nutrient acquisition capacity of *C. weissflogii* and potentially of diatoms more broadly.

In this study, we tracked *M. adhaerens* nitrogen transfer to *C. weissflogii*, as nitrogen can be readily traced using stable isotopes. Quantitative considerations suggest, however, that bacterial capture via spines alone is unlikely to fully satisfy the diatom’s nitrogen demands (Note S5): assuming complete transfer of bacterial nitrogen, intercepting approximately 88 bacteria per hour would be required to sustain diatom growth — a rate far exceeding both our experimental observations and encounter rate predictions (Fig. S4). Spine-mediated bacterial capture therefore likely acts synergistically with other nitrogen acquisition pathways in diatoms. Importantly, nitrogen may not be the most ecologically relevant currency of this interaction. Lysed bacteria are expected to release organic carbon, as well as a wide range of metabolites — including vitamins^5,69^, siderophores^70^, and growth-promoting infochemicals^6^ — that are frequently limiting for phytoplankton growth^5–7,70^ and may represent a significant ecological benefit of spine-associated bacterial interactions.

### Ecological significance and broader implications

Bacterial attachment to *C. weissflogii* spines is not restricted to *M. adhaerens*: four Vibrio strains and *Alteromonas macleodii* attached with comparable affinities. Together with previous observations of *Vibrio parahaemolyticus* attached to *C. weissflogii* spines^33^, *Bacteroides* associated with *Cytophaga* spp. spines^34^, and *Phaeobacter spp.* on *T. rotula* spines^35^, our findings show that spine attachment occurs across diverse diatom and bacterial taxa and suggest that individual diatoms can host multiple bacterial taxa on their spines. Whether bacterial attachment to diatom spines commonly translates into nutrient acquisition by the host across these diverse associations remains to be determined, but the recurrence of this interaction across multiple taxa raises the possibility that it constitutes a broader component of diatom–bacterium ecology. Establishing how widespread spine-mediated suspension feeding is among phytoplankton will require extending this experimental framework to additional bacterial and spine-bearing phytoplankton species and to natural communities. The answer will have potentially far-reaching consequences for our understanding of phytoplankton mixotrophy and marine nutrient cycling.

## MATERIAL AND METHODS

### Cell culturing and media preparation

*Conticribra weissflogii* (CCMP1051, Bigelow) was cultured in f/2 medium^71^ supplemented with 1.06×10⁻⁴ M silica (hereafter ‘f/2 + Si medium’). Cells were grown at 14°C on a 12:12 diurnal cycle at light intensity of 100 μmol photons m^-2^ s^-1^. Every 14 days, 1 mL of culture was transferred into 29 mL of fresh f/2 + Si medium.

*Marinobacter adhaerens* Hp15^36^ was grown in marine broth (Difco 2216, Sigma Aldrich L3022) at 27 °C shaken at 220 rpm, in 14 mL polypropylene round-bottom tubes (Greiner Bio-One 187262). eYFP-or DsRed-tagged derivatives^40^ were maintained with 100 µg mL⁻¹ ampicillin. Before every experiment, cultures were initiated from −80 °C glycerol stocks, grown overnight, and diluted 1:100 into fresh medium for a second overnight incubation to ensure active growth.

The same culturing conditions were applied to the following fluorescently labelled bacterial strains: *Alteromonas macleodii*-GFP (strain 27126, pLL104 plasmid, conjugated in this study as in ref), *Vibrio cyclitrophicus*-GFP (strain ZF270, pLL104 plasmid^72^, gift from the Cordero lab), *Vibrio coralliilyticus*-GFP (strain YB2, pLL104 plasmid^72^ conjugated in this study), *Vibrio ordalii* (strain 12B09, pGFP plasmid from Clontech laboratories^73^), and *Vibrio splendidus* (strain 13B01, pLL104 plasmid^72^, gift from the Cordero lab). All strains were maintained with 25 µg mL⁻¹ chloramphenicol to sustain fluorophore expression, except *V. ordalii*, for which spectinomycin resistance is encoded by the pGFP plasmid (100 µg mL⁻¹).

Artificial seawater (ASW) was prepared by dissolving Instant Ocean® Sea Salt in milliQ water, autoclaving, and filtering through a 0.2 µm membrane.

### Isolation of diatoms and attached bacteria from co-cultures

The diatom fraction (diatom cells and attached bacteria) was isolated from *C. weissflogii*–*M. adhaerens* co-cultures by filtering 2.5 mL of co-culture through an 8 µm Whatman® nitrocellulose membrane under minimal vacuum. Cells retained on the filter were washed with 20 mL of ASW to ensure complete removal of unattached bacteria and resuspended in 500 µL of ASW before observation.

### Growth assay

Diatom and bacterial densities in mono- and co-cultures were measured in triplicate at each time point by flow cytometry (see *Cytometry*).

Bacterial growth was assessed by measuring OD₆₀₀ over 48 h at 27°C in 96-well plates using a microplate reader (Synergy H1, BioTek), with 30 s orbital shaking before each measurement. Each well contained 200 µL of a 1:100 dilution of an overnight culture — itself initiated from a −80°C glycerol stock — in fresh marine broth, with eight replicates per condition.

### Microscopy

#### Epifluorescence microscopy

Samples were imaged on a Nikon Eclipse Ti2 inverted microscope equipped with a Prime BSI (PCIe) camera (Teledyne Photometrics), a 10X (NA 0.3), 20X (NA 0.45) and 40X (NA 0.6) objectives, and a Nikon Intensilight C-HGFIE pre-centered fiber illuminator. Fluorescence was detected using GFP (Semrock GFP-4050B), mCherry (Semrock m-Cherry-C), YFP (Nikon F46-003), or DAPI (Chroma 49028) filter sets.

#### Scanning electron microscopy (SEM)

7 mL and 3.5 mL of 8- and 18-day-old axenic *C. weissflogii* cultures (respectively) were centrifuged at 413 x g for 10 min and resuspended in 1 mL of supernatant. Concentrated diatoms (200 µL) were mixed with 300 µL of an overnight bacterial culture washed by centrifugation (1650 x g, 3 min) and resuspended in f/2 + Si medium. The final bacterial density was 10⁶ bacteria mL⁻¹. Co-cultures were incubated for 2 hours at 14°C. 200 µL of cocultures were then fixed with 1% (w/v) glutaraldehyde and deposited on hydrophilized 0.01% poly-L-lysine coated silicon. After 10 min incubation, the wafers were sequentially immersed for 5 min each in 2.5% glutaraldehyde solution (SOW, 27 practical salinity units, PSU), 1% osmium tetroxide, and SOW. Wafers were then passed through an ethanol drying series by sequential immersion in 0%, 30%, 50%, 70%, 90%, and 100% ethanol for 2 min at each stage, and finally three times in water-free ethanol. Wafers were dried using a critical point dryer with a cell monolayer protocol (CPD Tousimis, ETH Zurich Microscope Facility ScopeM) and were fixed to aluminum SEM stubs using silver paint. Samples were degassed for 24 h then sputter coated with 4 nm platinum-palladium (CCU-010 Metal Sputter Coater Safematic, ETH Zurich Microscope Facility ScopeM). SEM imaging was performed using an extreme high resolution (XHS) TFS Magellan microscope (ETH Zurich Microscope Facility ScopeM) with 50 pA current and an accelerating voltage of 5 kV for imaging and 15 kV for EDX. Scattered electrons were collected and imaged with both an immersion backscattered electron CBS detector and secondary electron TLD detector.

#### Confocal microscopy

Samples were examined using a LSM 780 (Zeiss) confocal microscope equipped with Airyscan detector (Carl Zeiss). FITC and diatom auto fluorescence were excited using laser lines of 488 and 561 nm wavelength, respectively. The detection windows for FITC and diatom auto fluorescence emission were set to 495–550 nm and long pass from 605 nm, respectively. Z-stack images were taken with a 63x (1.4 NA) oil objective. Images were collected and processed using the ZEN black software (Zeiss). *Cryo-electron tomography:*

Two samples were prepared. First, the diatom fraction — containing *C. weissflogii* cells and their spine-attached *bacteria* — was isolated from 14-day-old co-cultures initiated at 10⁴ diatoms mL⁻¹ and 5×10⁵ *M. adhaerens* cell mL⁻¹ (see *Isolation of diatoms and spine-attached bacteria from co-cultures*). Second, to image bacteria that had not been exposed to diatoms, 2 mL of an overnight *M. adhaerens* culture were centrifuged at 1650 x g for 3 min and resuspended in ASW.

Samples were mixed with 10 nm Protein A coated gold beads (Cytodiagnostics) at a 1:5 ratio and 3.5 µl of the sample was applied to glow-discharged copper EM grids (R2/1, Quantifoil), automatically blotted from the backside for 5–7 s and plunged into liquid ethane/propane^74^ using the Vitrobot Mark IV (Thermo Fisher Scientific).

Tilt series were collected on a Titan Krios G4 (Thermo Fisher Scientific) operating at 300 kV equipped with a BioContinuum imaging filter and a K3 direct electron detector (Gatan) using SerialEM^75^. After identification of targets by low magnification 2D screening, tilt series were acquired using a dose-symmetric tilt scheme, at a pixel size of 4.51 Å at the specimen level using 2° angular increments between tilts and a target defocus of −6 µm, covering an angular range of −60° to +60° and a total electron dose of 140–160 e^−^ Å^−2^.

Tilt series were drift-corrected using *alignframes* in IMOD^76^ and tomograms were reconstructed by weighted-back projection in IMOD at a binning factor of 4. To enhance the contrast, tomograms shown in this manuscript were CTF-deconvolved and filtered using *isonet*^77^.

### Staining

#### Diatom spines staining

Spines were visualized using the chitin stain calcofluor white (Sigma-Aldrich, 18909). 1 mL of diatom axenic cultures were incubated with 10 µg mL^-1^ calcofluor white in the dark at 4°C for 30 min. Stained cells were then observed using a Nikon Ti2 epifluorescence microscope mounted with a 40x and a 20x objective and a DAPI filter cube (Chroma 49028).

*Bacterial Viability Staining*:

Bacteria were incubated with either 1 µg mL^-1^ propidium iodide (ThermoFischer P3566) or 1.5 µg mL^-1^ Sytox Green (ThermoFischer S7020) for 15 minutes in the dark at 4°C. PI was used to assess the antibacterial effect of purified spines and fucoidan (Note S1); Sytox Green was used to quantify the proportion of lysed bacteria among spine-attached cells. Stained samples were either analysed using flow cytometry or epifluorescence microscopy (using a 20x objective).

#### Immunolabeling of diatom polysaccharides

*C. weissflogii* cells from 6-day axenic cultures were screened for four polysaccharide epitopes using epitope-specific monoclonal antibodies, following a previously established protocol^52^. The selection of these epitopes was based on prior reports highlighting the presence of the targeted polysaccharides in diatom extracellular polymeric substances^52,53^. The primary antibodies used in this study and their specificity can be found in Table S1. The secondary antibody was an anti-rat secondary antibody conjugated to FITC (F1763, Sigma Aldrich).

Briefly, diatom cultures were fixed with 1% formaldehyde and incubated for 1h at room temperature. Subsequently, 1 mL of each fixed culture was filtered at minimum vacuum intensity onto black polycarbonate filters (0.2 µm pore size, 47 mm diameter) and washed with 10 mL sterile Milli-Q water. Filters were air-dried on Whatman paper in Petri dishes for 20 min, then sectioned into pieces and single pieces were placed into 1.5 mL Eppendorf tubes. Samples were blocked with 400 µl of 5% PBS-MP (phosphate-buffered saline with 5% milk powder) for 1 h. After removing the blocking solution, 400 µl of primary antibody diluted 1:5 in PBS-MP, or 400 µl of PBS-MP alone for negative controls, was added and incubated for 1.5 h. Filters were washed four times with 1 mL PBS per wash. 300 µl of secondary antibody (anti-rat FITC-conjugated) diluted 1:100 in PBS-MP was added and incubated for 1.5 h in darkness. Post-incubation, filters were washed four times with PBS under dark conditions. Each filter piece was mounted on a 1 mm thick glass slide with embedding medium and covered with a coverslip. Prepared slides were stored at 4 °C.

Immunolabeled filters were examined with a confocal laser scanning microscope mounted with a 63x oil-immersion objective and an epifluorescent microscope mounted with a 40x objective.

#### Lectin staining

Diatom samples were prepared as described in the immunolabeling protocol. Filter sections were incubated with 300 µL of 5 µg mL^-1^ Aleuria aurantia lectin (AAL)-biotin conjugate –or 300 µl of PBS alone for negative controls– for 45 min at room temperature. Following incubation, filters were washed four times with 400 µL PBS to remove unbound lectin. Subsequently, filters were incubated with 2 µg mL^-1^ streptavidin-AlexaFluor 488 for 2 h in the dark. After incubation, filters were washed four times with PBS under dark conditions, mounted on 1 mm thick glass slides using embedding medium and stored at 4 °C. Lectin fluorescence was assessed using epifluorescence microscopy.

### Cytometry

Diatom and bacterial cell counts were measured using a cytometer (CytoFLEX B3-R2-V2, Beckman Coulter) equipped with a 488 nm laser. Diatom abundance was quantified using forward scatter and autofluorescence (B585 channel). Bacterial abundance was quantified using forward scatter and either red fluorescence of DsRed-labelled strains and Propidium Iodide-positive bacteria (B585 channel) or yellow/green fluorescence of SYBR Green, YFP- and GFP-labelled strains (B525 channel). Flow cytometry data were acquired and analyzed using CytExpert software (Beckman Coulter).

### Raman resonance spectroscopy

Nitrogen was chosen as a marker of metabolic exchange because it is a key limiting nutrient for phytoplankton^78^ and its isotopes are experimentally tractable and well-suited for isotope-probing studies^79,80^. *M. adhaerens* was grown overnight in marine broth (see *Cell culturing*), then diluted 1:100 into artificial seawater supplemented with 1g.L^-1^ ^15^N-labeled Celtone (Cambridge Isotope Laboratories) and incubated for 48 hr. 2 mL of cells were washed twice with artificial sea water (centrifugation at 1650 x g for 3 min) and resuspended in 1 mL of ASW. 200 µL of this suspension were stained with SYBR Green YBR Green (5 μM final concentration, Sigma Aldrich CAS 163795-75-3), and bacterial density measured using flow cytometry (see *Cytometry*). The resulting bacterial suspension was diluted in 10 mL of f/2 + Si medium devoid of nitrate (prepared omitting NaNO₃) to achieve a final concentration of approximately 5×10^4^ cells mL^-1^. One mL of *C. weissflogii* cells from a 14-day-old axenic culture was added to the final bacterial suspension, and cocultures were incubated at 14°C for 6 days (see *Cell culturing and media preparation*). Control cultures of, *C. weissflogii* were grown axenically in regular f/2 + Si medium (negative control, “^14^N-diatoms”) and in f/2 + Si medium prepared with ^15^N-nitrate instead of ^14^N-nitrate (positive control,”^15^N-diatoms”).

Raman measurements were performed using a confocal Raman microspectroscope (LabRAM HR Evolution, Horiba Scientific, France). The system comprises an upright microscope (BxFM, Olympus) integrated with components for Raman analysis, including a continuous wave Nd:YAG laser at 785 nm (10 mW), whose excitation generates data specifically reflecting chlorophyll and carotenoids pigments^43–45^, a 60x objective (MPlan N, Olympus), a 300 grooves/mm grating blazed at 6600 nm, and a back-illuminated deep-depleted CCD detector. Spectra of individual diatom cells were acquired within the 600–1600 cm⁻¹ range, covering most cellular signals, with a 1-second laser exposure and a 100 µm confocal pinhole. Prior to acquisition, 5 µl of samples were loaded into a 375 µm high microfluidic chamber. For each diatom, three Raman spectra were recorded, targeting different locations of the cell to obtain an average signal and minimise the potential effect of intracellular heterogeneity. Colonization status of diatoms was examined prior to each Raman measurement using a 60x air objective. Diatom cells with bacteria attached to the cell wall, rather than to a spine or the base of a spine, were excluded from the analysis. Raman measurements were taken a couple of micrometers away from the spine-attached bacteria, confirming that the peak downshift arose from diatom metabolism rather than interference from the bacteria.

Diatom Raman spectra were processed using a custom MATLAB ® (Mathworks 2026a) script. Briefly, the data underwent smoothing (de-noising) and baseline subtraction using the Savitzky–Golay filter (with polynomial order 3 and frame length 7) and a polynomial-based algorithm (2^nd^ order polynomial), respectively. Peaks in Raman spectra were identified using the MATLAB *peakfit()* function with a window parameter of 10.

Comparison of the mean Raman spectra of diatoms grown with ^14^N and ^15^N revealed four peaks (at 916, 1222, 1283 and 1375 cm^-1^, Fig. S3) exhibiting significant downshift along with ^15^N incorporation that correspond to nitrogen-containing moieties in chlorophyll^43–45^. Among these peaks, the one centered at 916 cm^-1^ showed the largest downshit along with ^15^N incorporation and was therefore selected as a proxy for ^15^N incorporation (Fig. S3). For the analysis of colonized and uncolonized diatom cells from cocultures with ^15^N-labeled *M. adhaerens*, the initial peak position was set to 916 cm^-1^. The mean peak position was first computed across all measurements within each experiment, and *peakfit()* was then reapplied using this experiment-specific mean value as the initial peak position.

### Microfluidic assay

To investigate the microscale dynamics of bacterial and beads attachment to *C. weissflogii*, a custom-designed microfluidic device was used (Fig. S8). The device comprised a single channel (height: 106 µm; width: 595 µm) containing six pillar blocks, each consisting of four lines of 30 µm-diameter micropillars. Lines within each block were spaced 150 µm apart, with inter-pillar spacings of 70, 60, 50, 40, 30, and 20 µm, respectively. The chip was fabricated using standard soft lithography in polydimethylsiloxane (PDMS) containing 10% (w/w) cross-linking agent (Sylgard 184 Silicone Elastomer Kit, Dow Corning) and bonded to a 75 mm × 25 mm × 1 mm glass slide using a plasma oven (model *Zepto*, diener electronic).

*C. weissflogii* cells were harvested from 6- or 14-day-old axenic cultures grown in f/2 + Si medium by centrifugation of 5 mL at 413 x g for 10 min. After removing 4.5 mL of the supernatant, the pellet was gently resuspended in the remaining 500 µL. This suspension was introduced into the microfluidic channel using a 1 mL BD plastic syringe.

Bacterial and suspensions were prepared from overnight cultures of *M. adhaerens*-eYFP, *A. macleodii*-GFP, *V. coralliilyticus*-GFP, *V. ordalii*-GFP, *V. splendidus-GFP*, and *V. cyclitrophicus-GFP* by centrifugation of 2 mL culture at 1650 x g for 3 min. Pelleted cells were resuspended in 1 mL artificial seawater and diluted to a final density of 5×10⁶ cells mL^-1^ in a total volume of 2 mL artificial seawater. For experiments involving lysed bacteria, cell suspensions were heat-shocked at 80 °C for 15 min and then kept on ice for 5 min prior to loading. Successful lysis was confirmed by flow cytometry using propidium iodide (1 µg mL⁻¹), which was also used to track bacteria in the microfluidic device.

For experiments with beads, yellow/green fluorescent raw polystyrene and carboxylated polystyrene beads (ThermoFisher, F13081 and F8823) were diluted to 5×10⁶ beads mL^-1^ in a total volume of 2 mL ASW.

The bacterial (or bead) suspension was loaded into a 1 mL BD plastic syringe connected to the microfluidic channel inlet, taking care to avoid bubble formation. Bacteria (or beads) were introduced into the channel at a flow rate of 5 µL h^-1^ using a syringe pump (Harvard Apparatus 11 Pico Plus Elite Syringe Pump), corresponding to a flow velocity similar to *Marinobacter* swimming speed^47^ (20 µm s^-1^). The effective flow speed in the microfluidic channel was measured by tracking fluorescent beads to ensure it corresponds to the value imposed by the microfluidic pump (Note S3).

Because individual diatom spines are poorly resolved by epifluorescence and light microscopy, attachment could not be quantified at the level of individual spines and was therefore scored per diatom cell. An attachment event was defined as the immobilization of a bacterium or a bead adjacent to a diatom cell. Once attached, bacteria and beads remained associated with the diatom, exhibiting only small displacements that matched the residual movement of the diatom cell.

### Bacterial binding assay

To assess bacterial binding affinity to commercially available fucoidan (from *Fucus vesiculosus*, Sigma-Aldrich CAS-9072-19-9) and chitin (Sigma-Aldrich CAS-1398-61-4) we employed a high-throughput binding assay. Wells of a flat-bottom 96-well plate (Thermo Fisher Scientific, 167008) were filled with 200 µL of each test compound at 1 g L^-1^ in Milli-Q water (n = 8 wells per treatment). Milli-Q water alone was used as a negative control. Plates were incubated overnight at 30 °C without lids to allow complete solvent evaporation and surface adsorption of the compounds. The following day, 200 µL of an HP15- eYFP bacterial suspension (10⁶ cells mL^-1^ in artificial seawater) was added to each well. Plates were sealed with parafilm and incubated overnight at 30 °C under static conditions. On day three, the supernatant from each well was carefully removed using a multichannel pipette. Wells were then gently washed three times with 200 µL of artificial seawater to remove unbound cells, taking care not to disturb surface-bound biofilms. A final 10 µL of artificial seawater was added to each well to prevent drying before imaging. Epifluorescence microscopy was performed using a 10× objective, and bacterial attachment was quantified with a custom ImageJ macro. On fucoidan-coated wells, attached *M. adhaerens* formed dense biofilms that precluded reliable counting of individual cells. Bacterial attachment was therefore quantified as the fraction of the imaged well surface covered by bacteria. The same analysis was applied to uncoated wells used as negative controls. On chitin-coated wells, bacterial attachment was substantially lower, and the chitin coating generated heterogeneous background fluorescence that prevented reliable surface coverage measurements. Attachment was therefore quantified by counting the number of bacteria per unit surface area.

### *C. weissflogii* spines purification

To obtain a pure and concentrated suspension of diatom spines, cultures were subjected to a sequential centrifugation-based purification protocol. An aliquot of 5 mL of 6-day axenic diatom culture was centrifuged at 1000 x g for 30 min at room temperature to pellet intact diatom cells. Following centrifugation, 4.5 mL of the supernatant was carefully collected without disturbing the cell pellet and transferred to a new tube. The collected supernatant was centrifuged at 20000 x g for 30 min at room temperature to pellet the spines. The resulting pellet was gently resuspended in 500 μL of filtered artificial seawater to wash the spines. The suspension was centrifuged again at 20000 x g for 30 min at room temperature, and the pellet was resuspended in 500 μL of filtered artificial seawater for a second wash. The final suspension was examined by light microscopy to confirm the absence of diatom cell bodies. Spine concentration was then determined using a hemocytometer under a light microscope.

### Statistics

All analyses were conducted in Python (3.12) using *SciPy* and *Statsmodels*.

#### Beads data

Effects of bead-type, age of diatom cultures and their interaction on the mean number of beads per diatom were assessed using two-way ANOVA with biological replicates treated as independent observations. Model assumptions were evaluated using quantile–quantile plots to assess normality and Levene’s test for homogeneity of variance.

Time series of bead accumulation were analyzed using a segmented regression framework to identify potential changes in beads accumulation rate. For each dataset, a candidate breakpoint (*t_b_*) was tested at all interior time points (excluding a margin of 5 time points from each boundary to ensure stable estimation). At each candidate position, a hinge function model y=β_0_+β_1_t+β_2_(t−t_b_)_+_ was fitted using ordinary least squares, where (t−t_b_)_+_=max(0, t−t_b_). The optimal breakpoint was defined as the time index minimizing the Sum of Squared Errors (SSE) across all candidate fits. This procedure was applied independently to each bead type. Analysis of SSE revealed a clear breakpoint at ∼10 minutes for the carboxylated beads (Fig. S7), whereas no sharp transition was observed for the unmodified polystyrene beads (Fig. S7). Following breakpoint identification, a continuous segmented regression model was fitted using the selected breakpoint to estimate pre- and post-break colonization rates.

#### Growth assay

Outliers within technical replicates for each treatment were identified and excluded based on Grubbs’ test. Differences among treatments were assessed using one-way analysis of variance (ANOVA) followed by Tukey’s honestly significant difference post-hoc test for the experiment with fucoidan (Note S1) and with an independent two-sample t-test for the experiment with purified spines (Note S1). Assumptions of normality and homogeneity of variances were evaluated using the Shapiro–Wilk and Levene’s tests, respectively.

#### Bacterial viability assay

The effect of fucoidan or purified spines (Note S1) on the proportion of PI-positive bacteria at different times post incubation was analyzed using a two-way ANOCA with biological replicates treated as independent observations. Assumptions of normality and homogeneity of variances were evaluated using the Shapiro–Wilk and Levene’s tests, respectively.

#### Raman spectral analysis

Raman shifts of uncolonized and colonized diatoms were obtained from three independent experiments, corresponding to three independent cocultures. To account for baseline shifts in Raman peak position between experiments, the primary analysis was performed on experiment-level contrasts. Within each experiment, we computed the mean position of the 916 cm^-1^ peak separately for uncolonized and colonized diatoms and defined the contrast *d_j_* = *Y_u,j_* – *Y_c,j_* where *Y_u,j_* and *Y_c,j_*denote the mean 916 cm^-1^ Raman peak position among uncolonized and colonized diatoms, respectively, in experiment *j*. Positive values of *d_j_* therefore indicate higher Raman peak positions in uncolonised diatoms relative to colonised diatoms from the same experiment. The primary estimand was the mean experiment-level contrast, 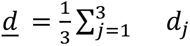 representing the average uncolonized-minus-colonized Raman shift difference across independent experiments. Because the hypothesis was directional, namely that colonized diatoms exhibit a lower 916 cm^-1^ Raman peak position than uncolonized diatoms, the three experiment-level contrasts were tested against zero using a one-sided one-sample t-test: *H*_0_ : *<u>d</u>* ≤ 0, *H*_1_ : *<u>d</u>* > 0.

Because the primary analysis was based on only three independent experiment-level contrasts (two degrees of freedom), we also performed an observation-level sensitivity analysis using all individual-diatom Raman shift measurements. We fit an ordinary linear regression model with colonisation status and experiment as categorical predictors: *Y_ij_* = *α* + *βG_ij_* + *γ_j_* + *ε_ij_*, where *Y_ij_* denotes the 916 cm^-1^ Raman peak position for diatom *i* in experiment *j*, *G_ij_* is an indicator for colonisation status, and *γ_j_* represents experiment-specific baseline differences in Raman peak position. Colonisation status is encoded as *G_ij_*=0 for uncolonised diatoms and *G_ij_*=1 for colonised diatoms, so that *β* estimates the colonised-minus-uncolonised difference. Thus, negative values of *β* indicate lower Raman shifts in colonised diatoms.

The coefficient estimate in this sensitivity analysis was the ordinary least-squares estimate, but inference for the colonization-status contrast used an HC3 heteroskedasticity-consistent standard error. HC3 estimates coefficient uncertainty using leverage-adjusted squared residuals rather than assuming a common residual variance and is appropriate here because the observation-level analysis involved modest and unequal sample sizes across colonization-status-by-experiment groups together with evidence of unequal Raman-shift variability^46^. This observation-level analysis was used to assess whether the estimated effect was consistent with the primary experiment-level contrast analysis while adjusting for experiment and allowing unequal variances across groups.

#### Binding assay

Differences in bacterial attachment density between control and treatment conditions were assessed using the Shapiro-Wilk test for normality and Levene’s test for equality of variances. For the fucoidan experiment, both assumptions were satisfied and a two-sample Student’s t-test was applied. For the chitin experiment, the normality assumption was violated, and Welch’s t-test was used instead.

## Acknowledgements

We gratefully acknowledge funding from a Gordon and Betty Moore Foundation Symbiosis in Aquatic Systems Initiative Investigator Award (GBMF9197; https://doi.org/10.37807/GBMF9197), the Simons Foundation (through the Principles of Microbial Ecosystems collaboration; grant 542395FY22), the Swiss National Science Foundation (grant 205321_207488 and Sinergia grant CRSII5-186422) and NCCR Microbiomes (a National Centre of Competence in Research consortium funded by the Swiss National Science Foundation; grant numbers 51NF40_180575 and 51NF40_ 225148) to R.S. We thank K. S. Lee for his help with setting up the Raman experiment and analyzing the results. We thank R. Naisbit for scientific editing.

---

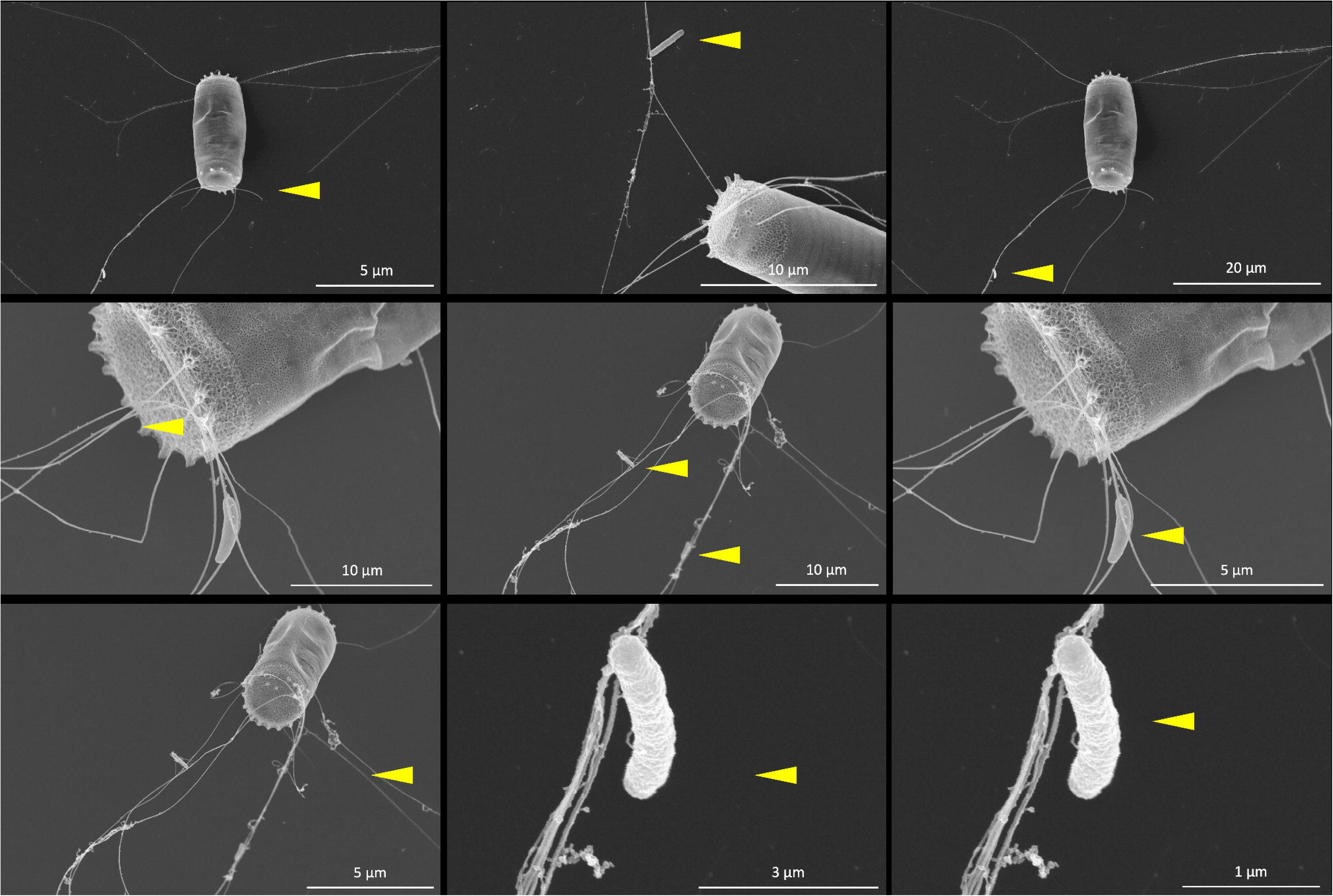

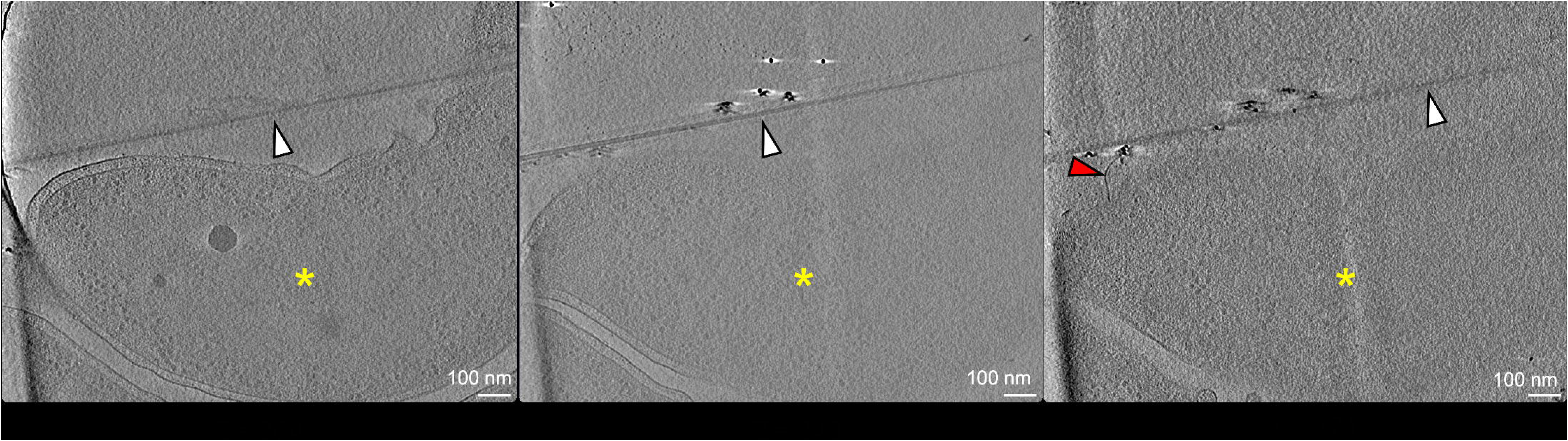

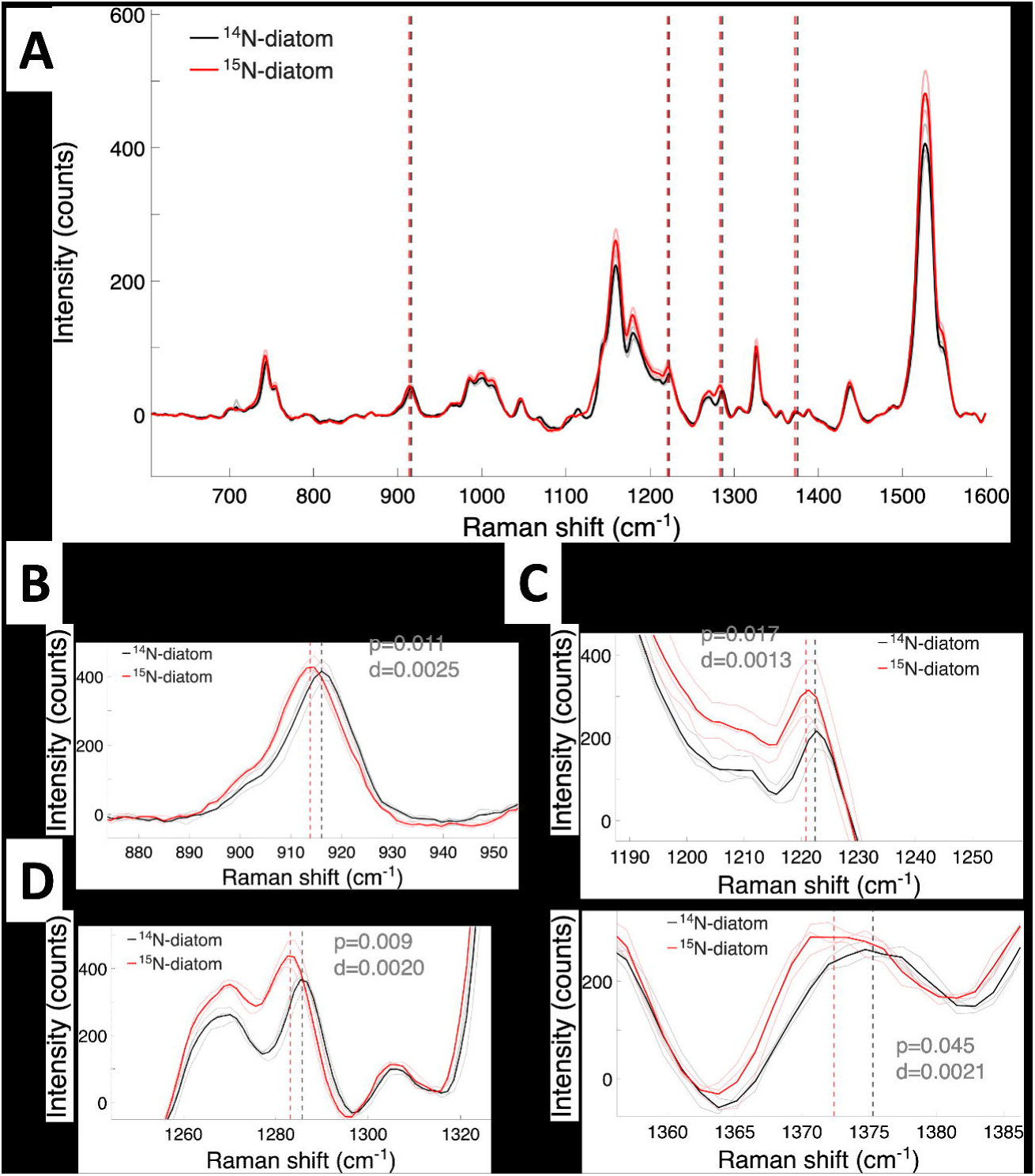

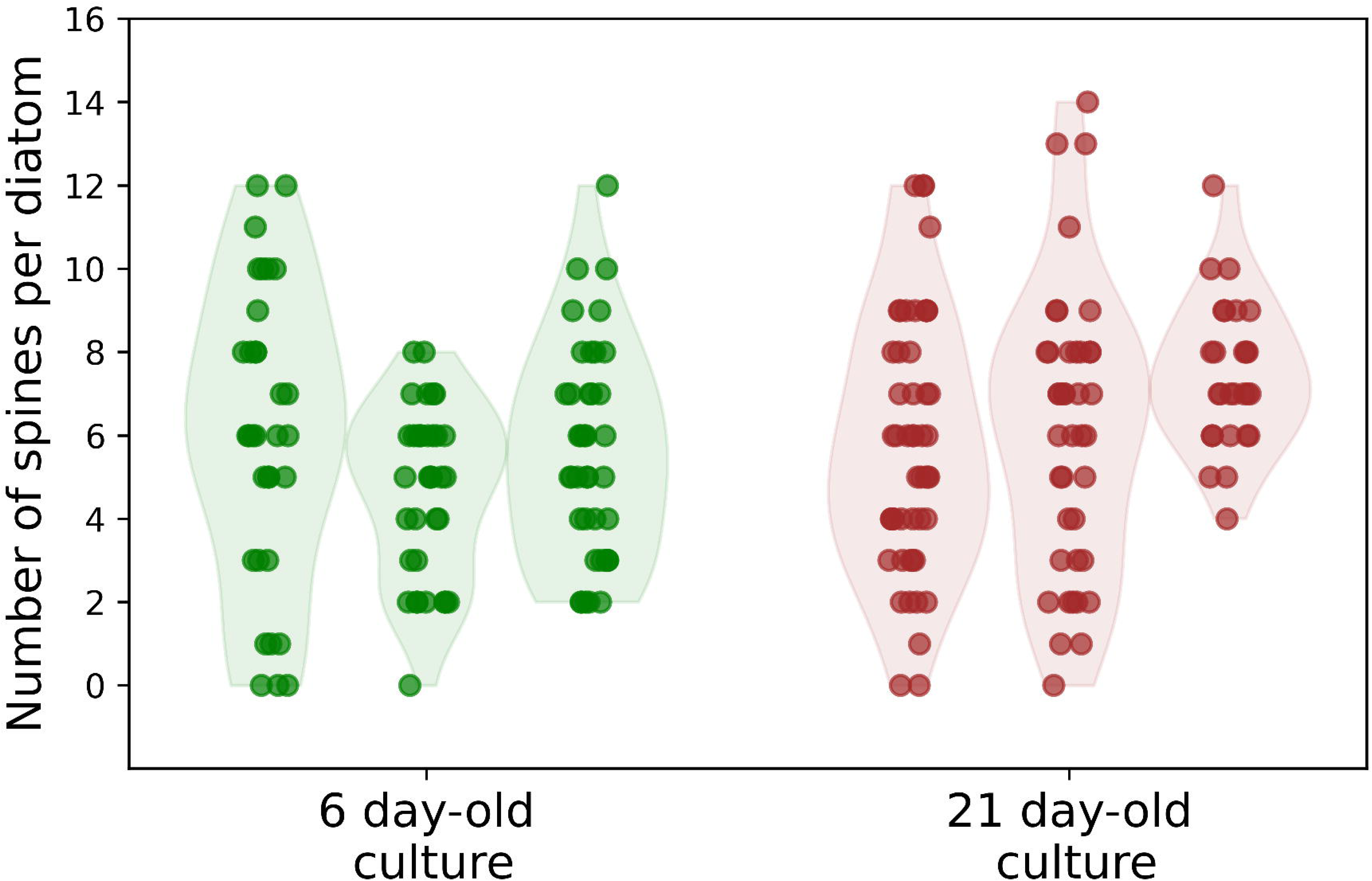

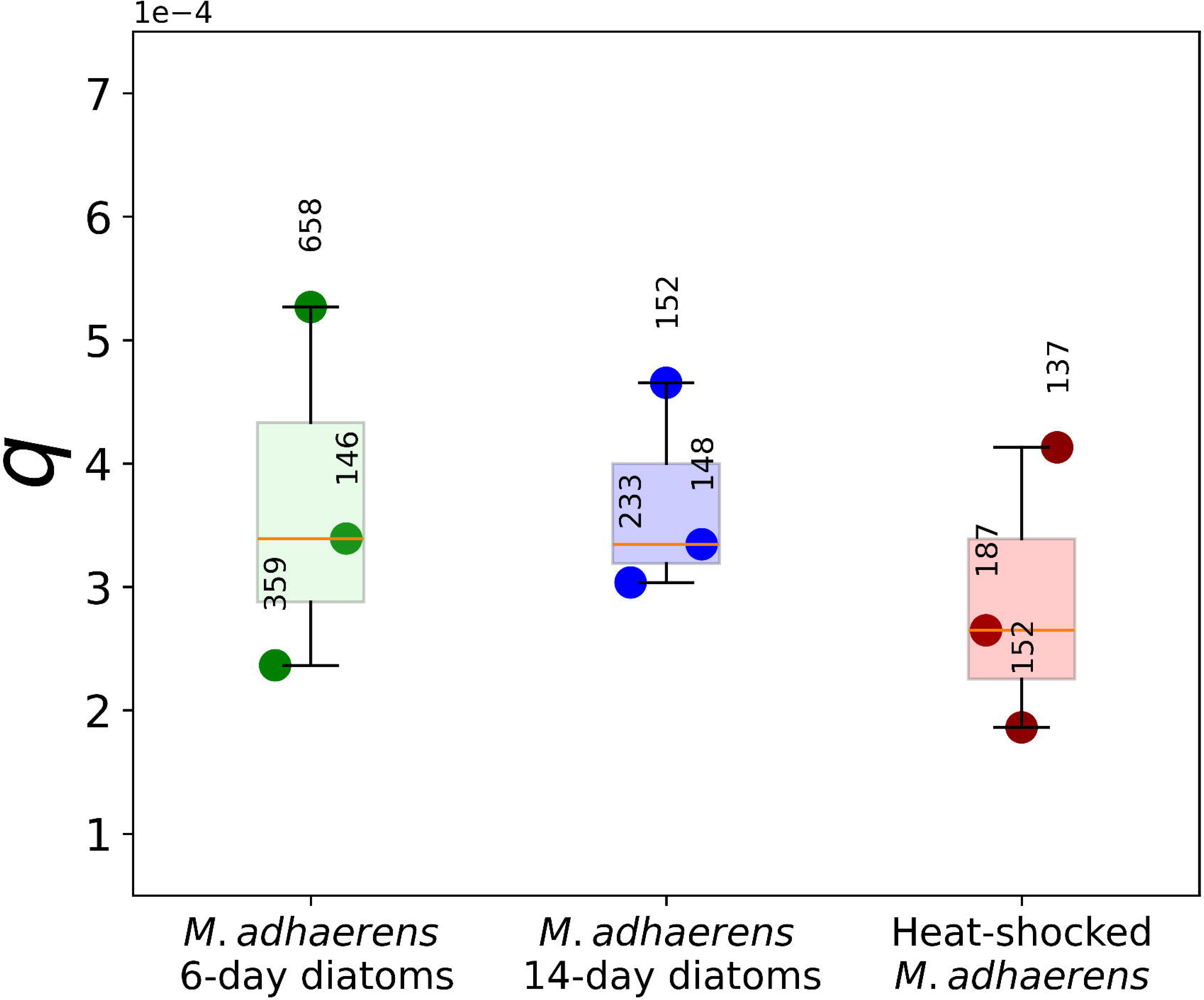

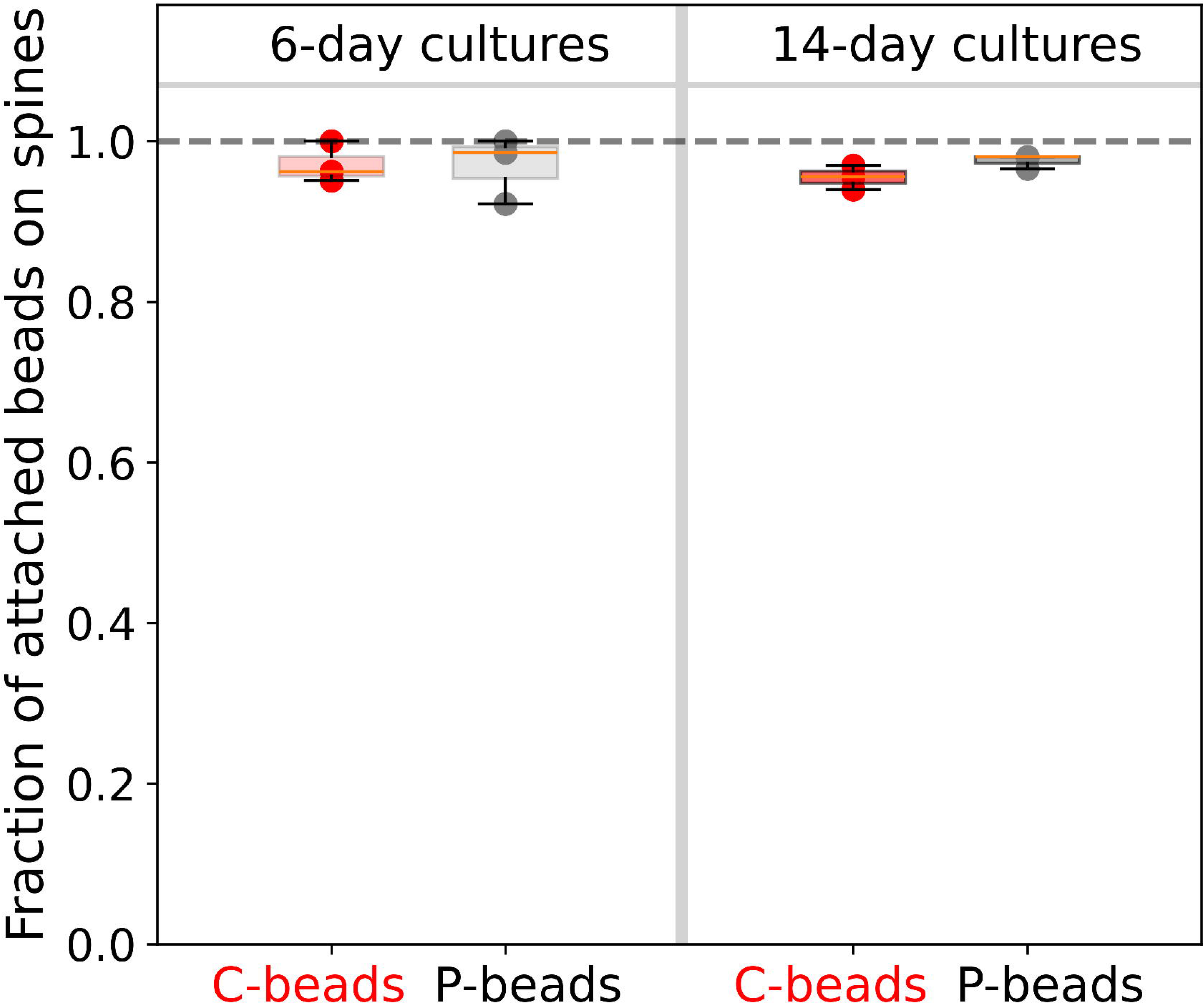

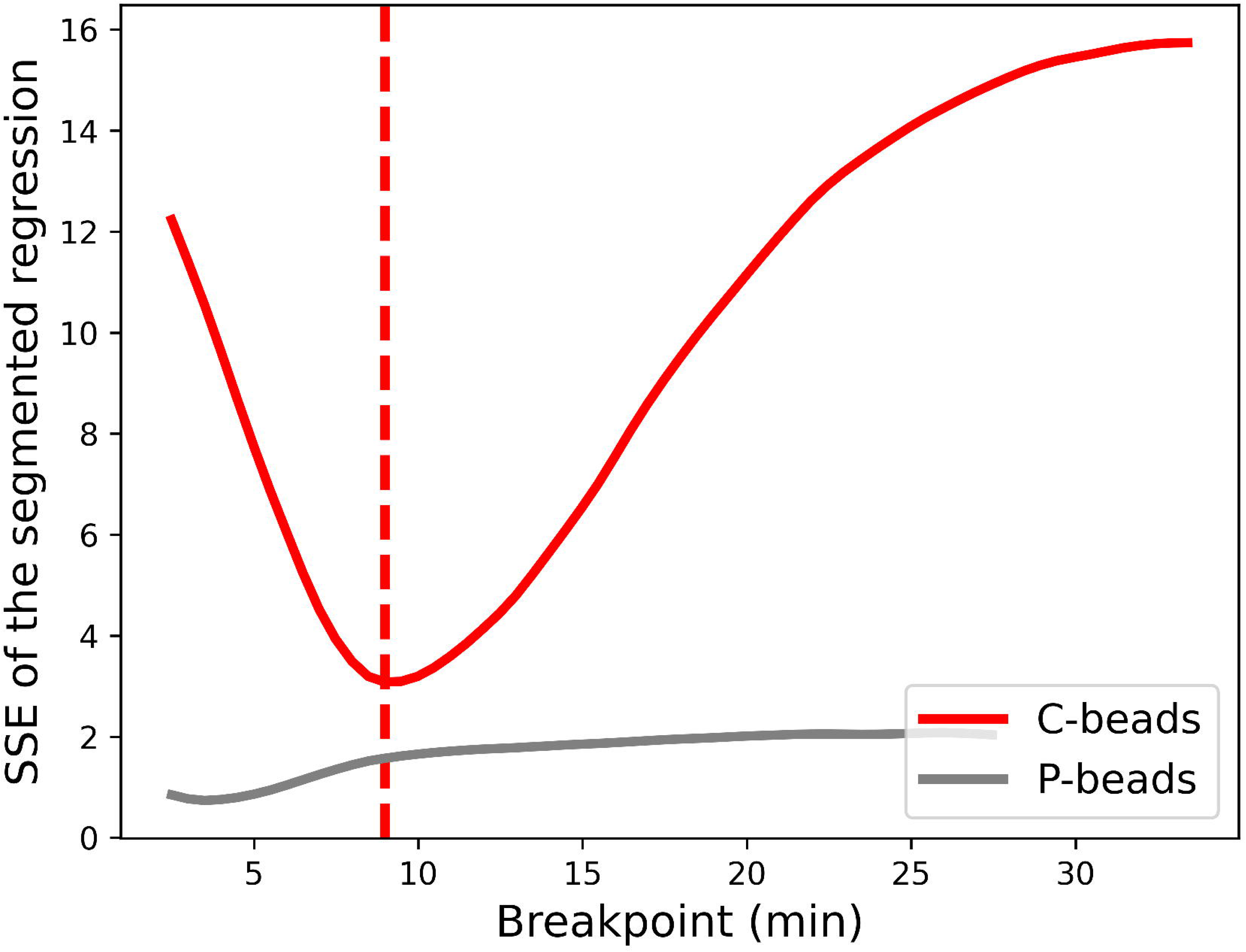

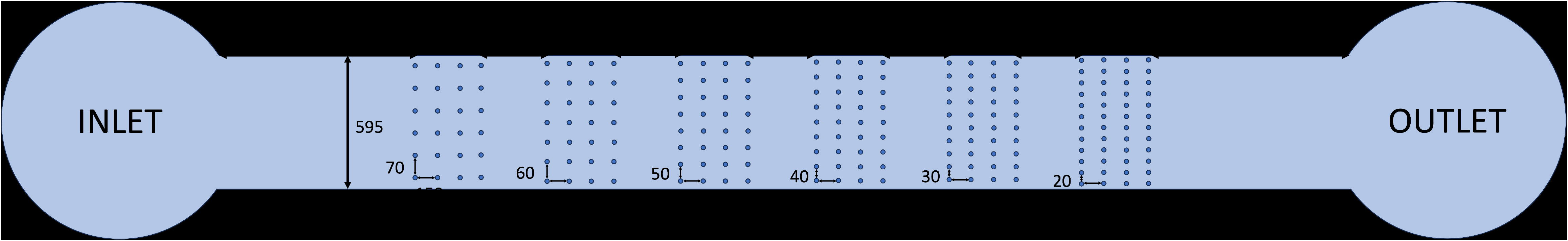

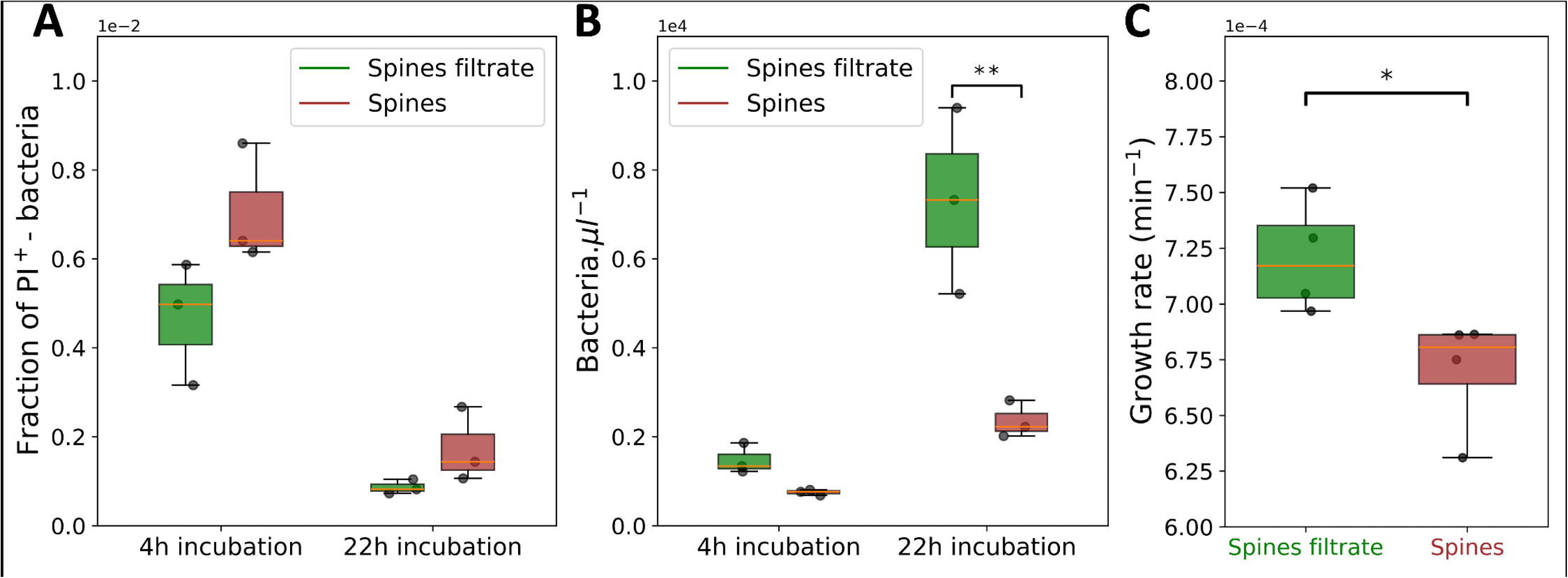

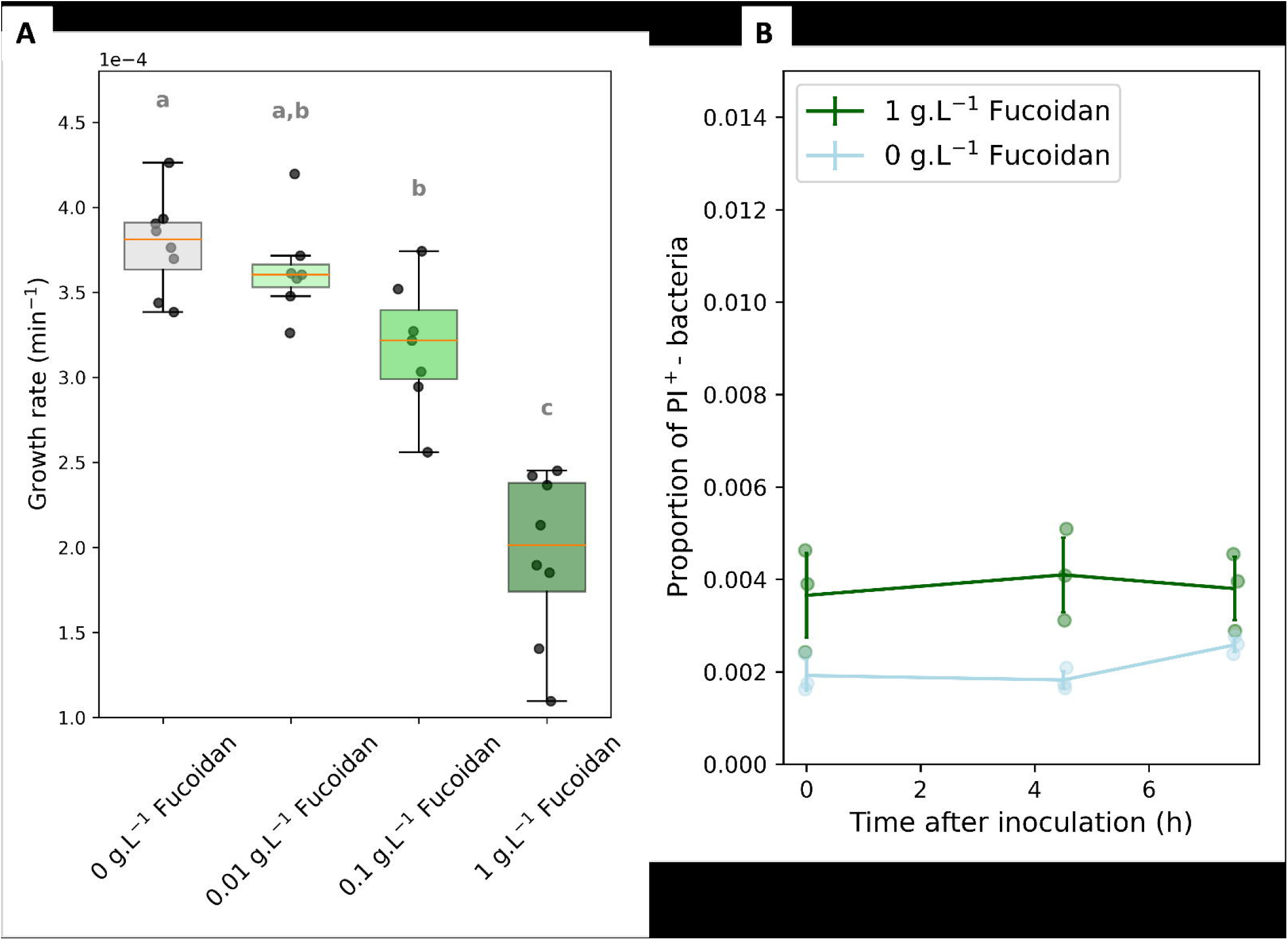

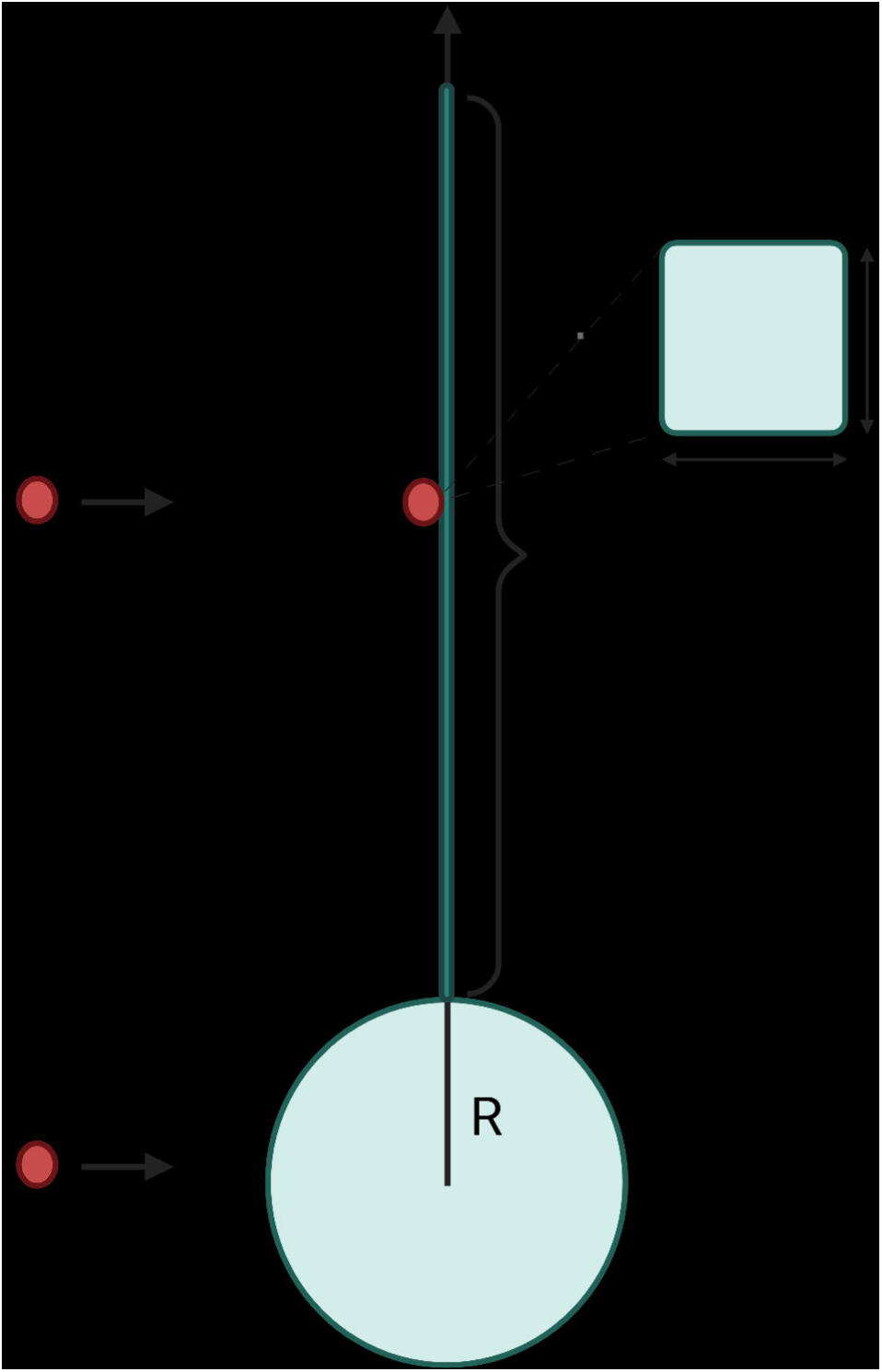

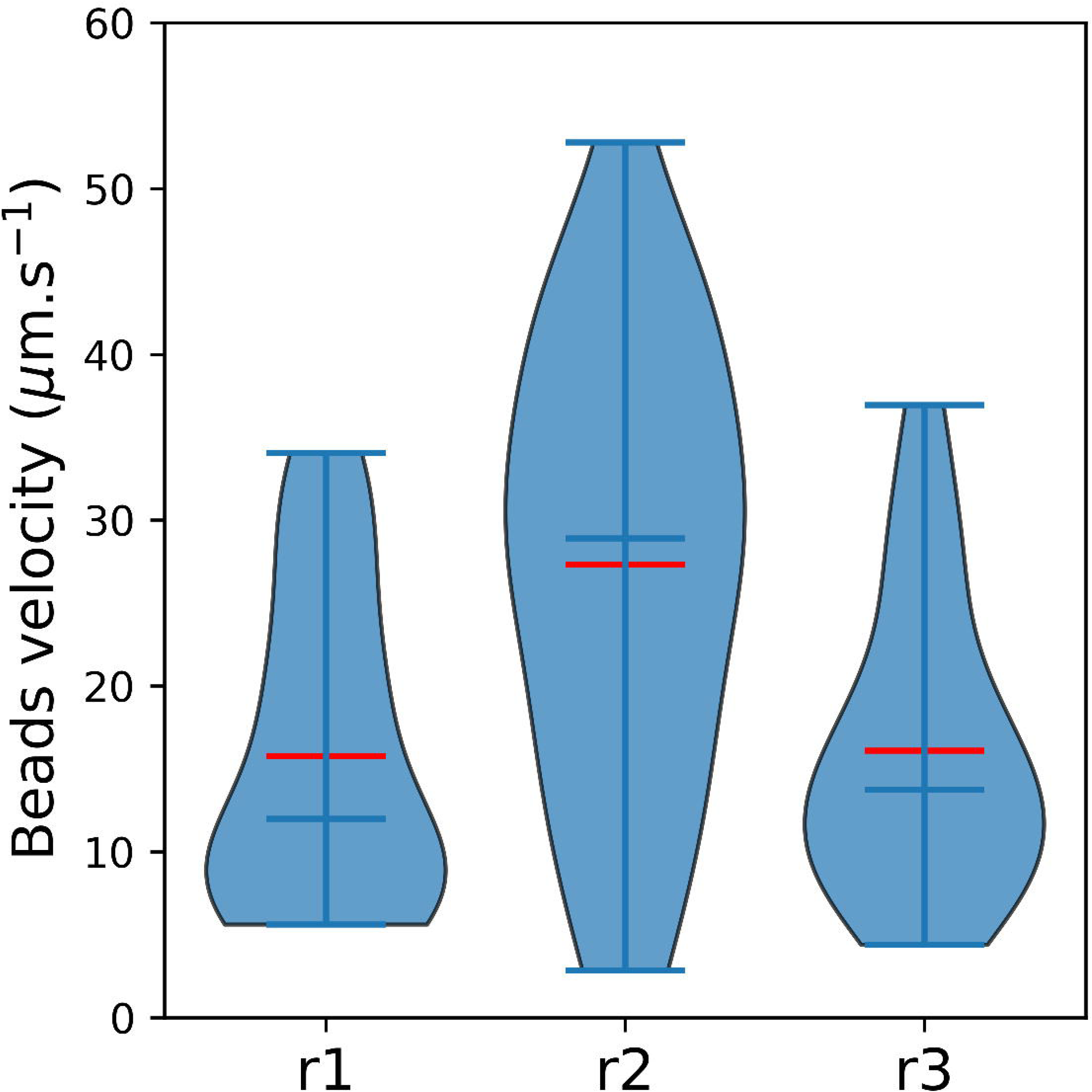

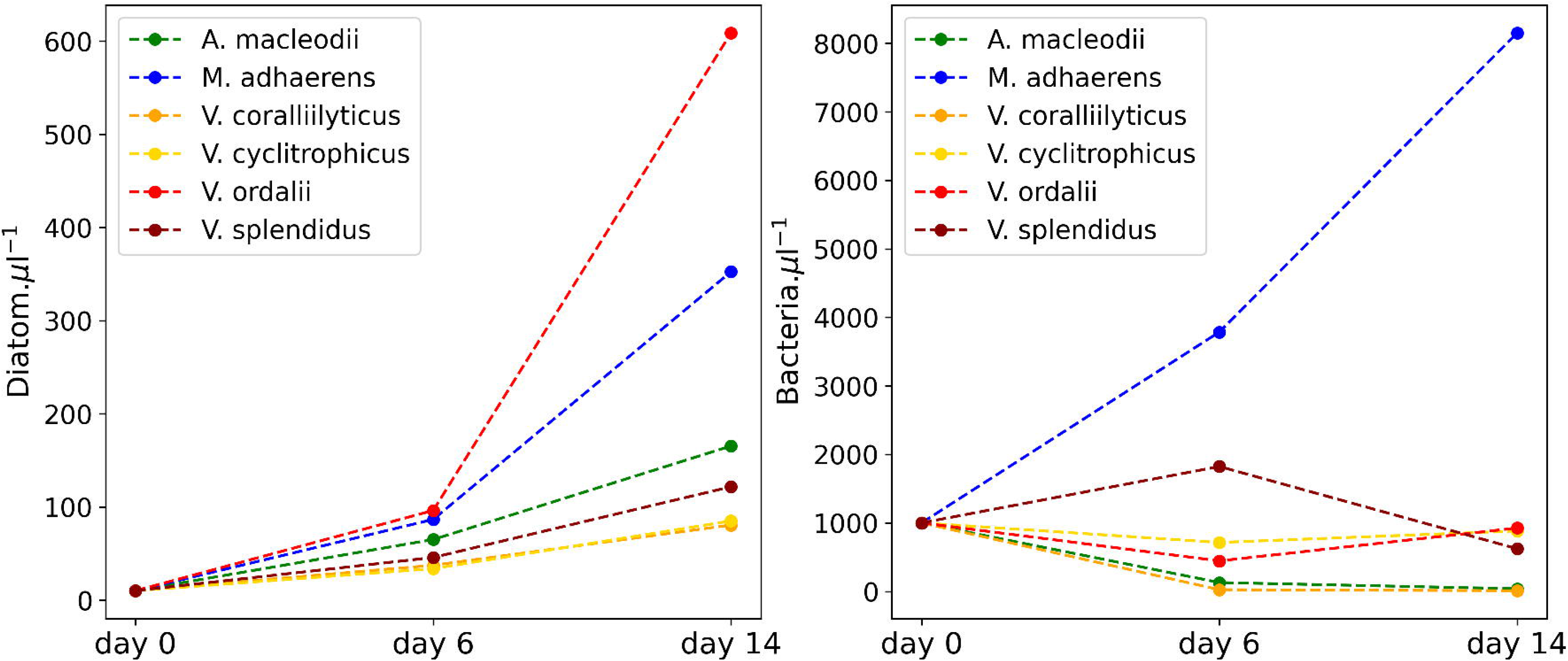

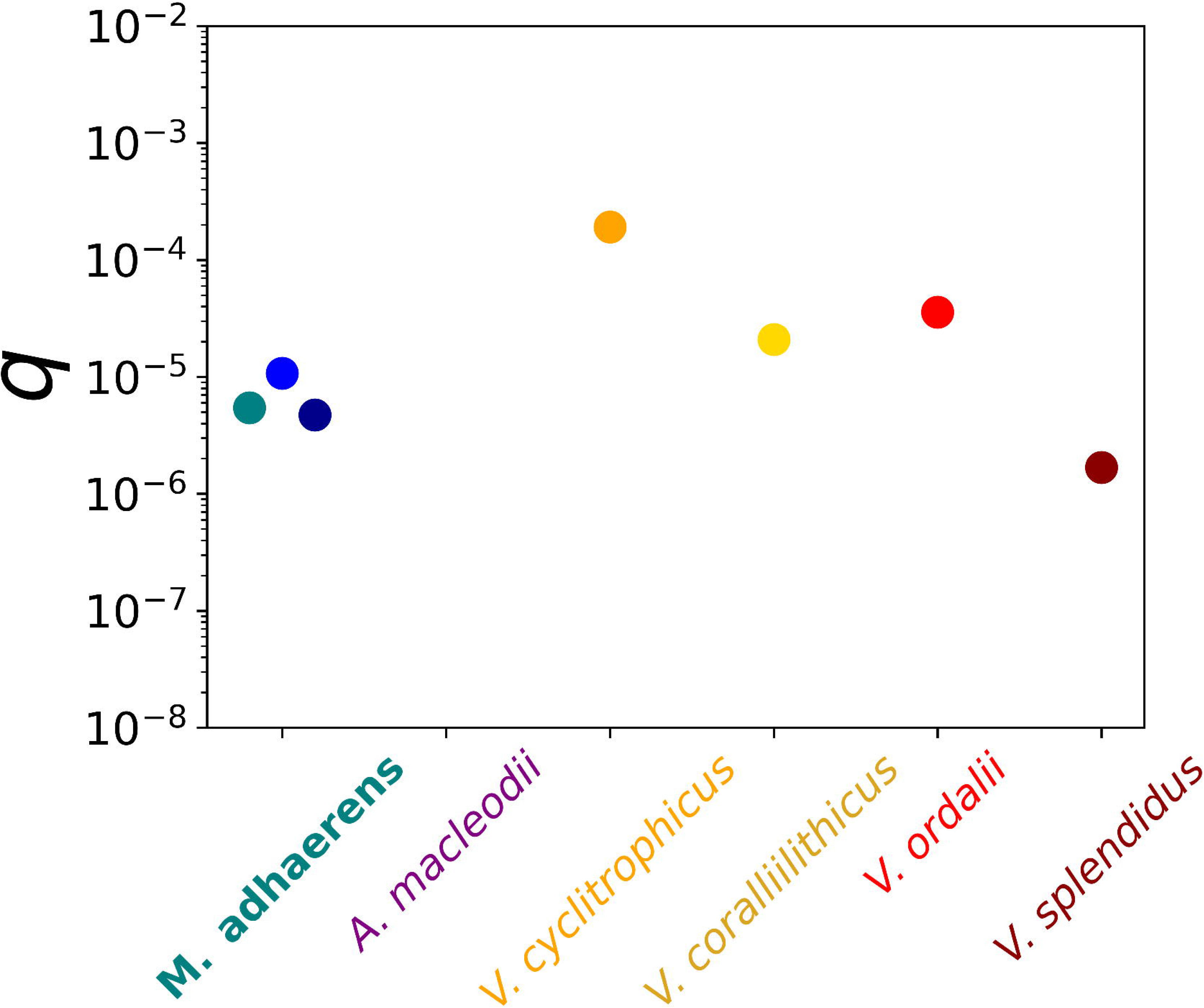

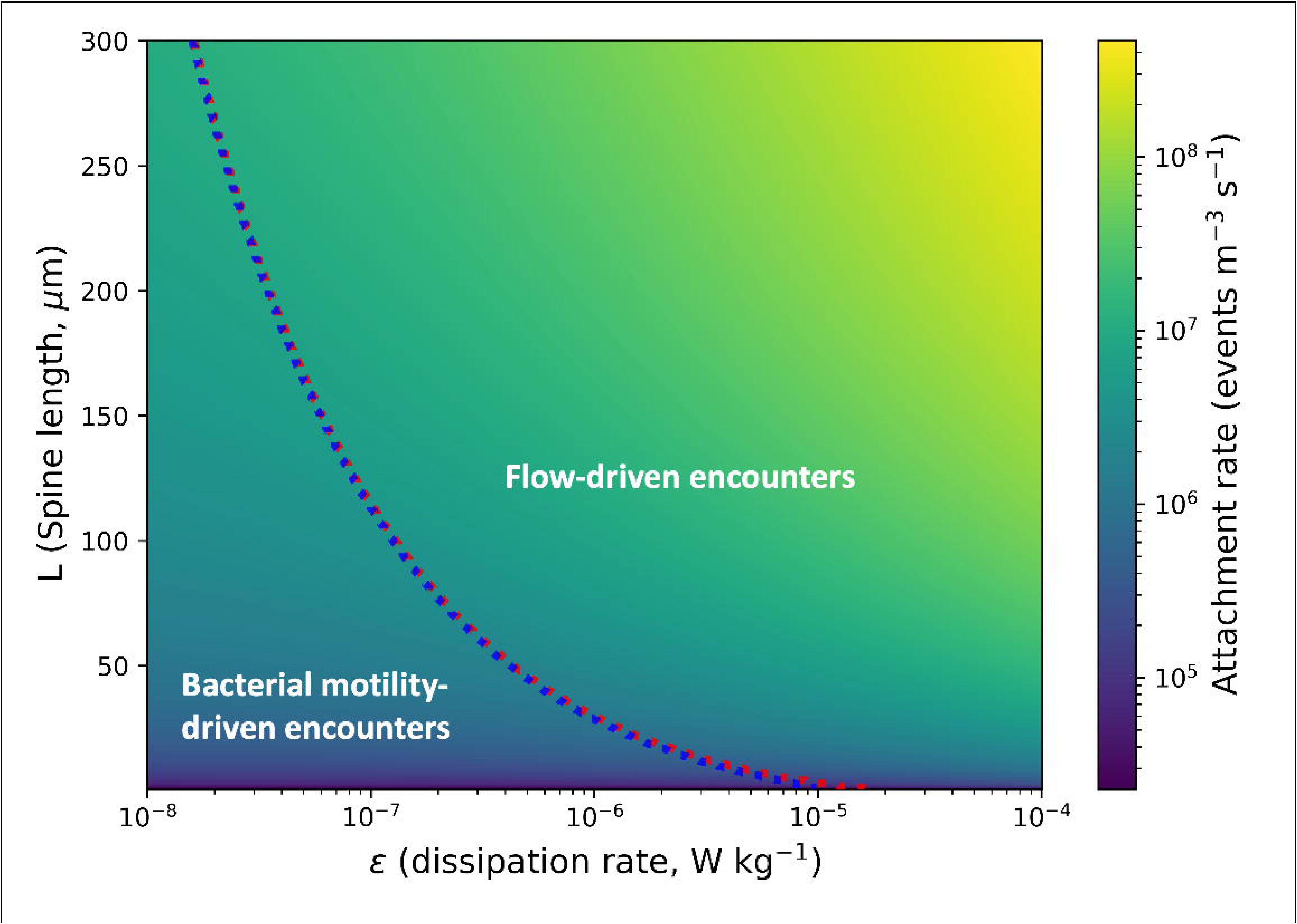

