## Supplementary Files for "Bacterial capture and lysis on diatom spines reveal a suspension-feeding strategy"

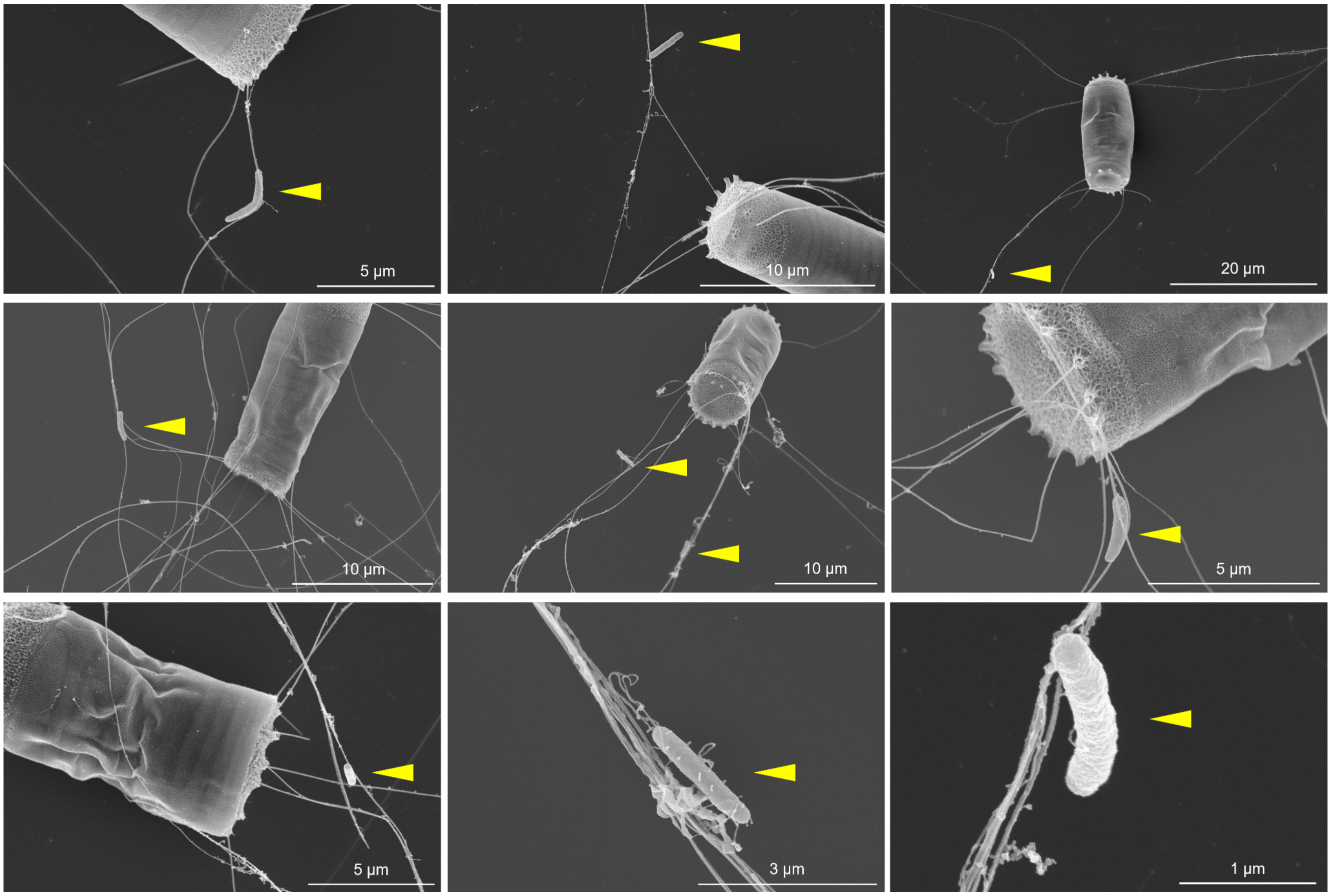


**Figure S1 | *M. adhaerens* attaches to *C. weissflogii* spines.**

SEM images of *M. adhaerens* (yellow arrowhead) interacting with the spines of *C. weissflogii.*

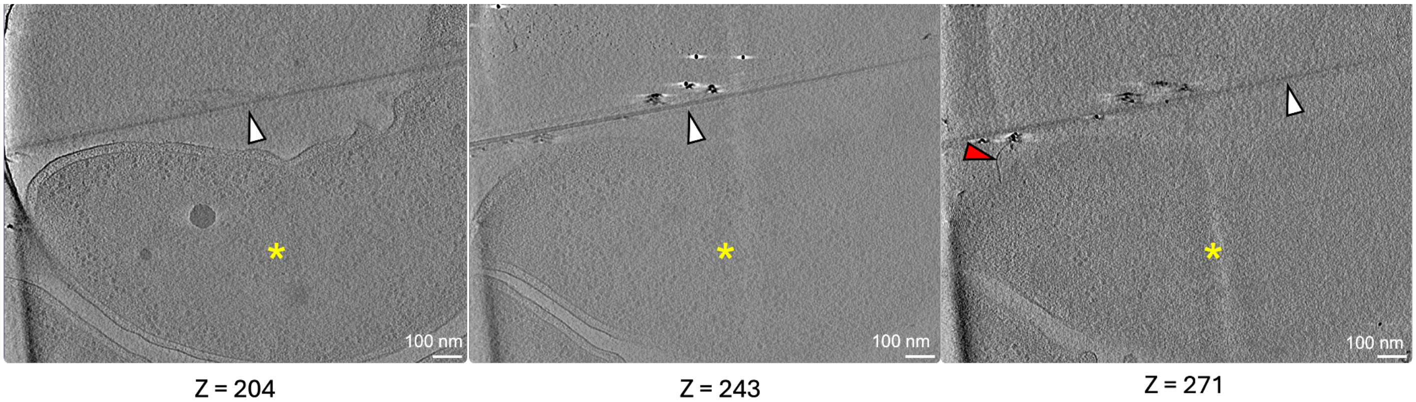


**Figure S2 | *M. adhaerens* interacting with *C. weissflogii* via a pilus.** Three tomographic slices at different depths through the same volume show *M. adhaerens* (yellow asterisc) interacting with *C. weissflogii* spines (white arrowhead). A bacterial pilus (red arrowhead) extends from the cell surface and contacts the spine, possibly suggesting a role in mediating attachment.

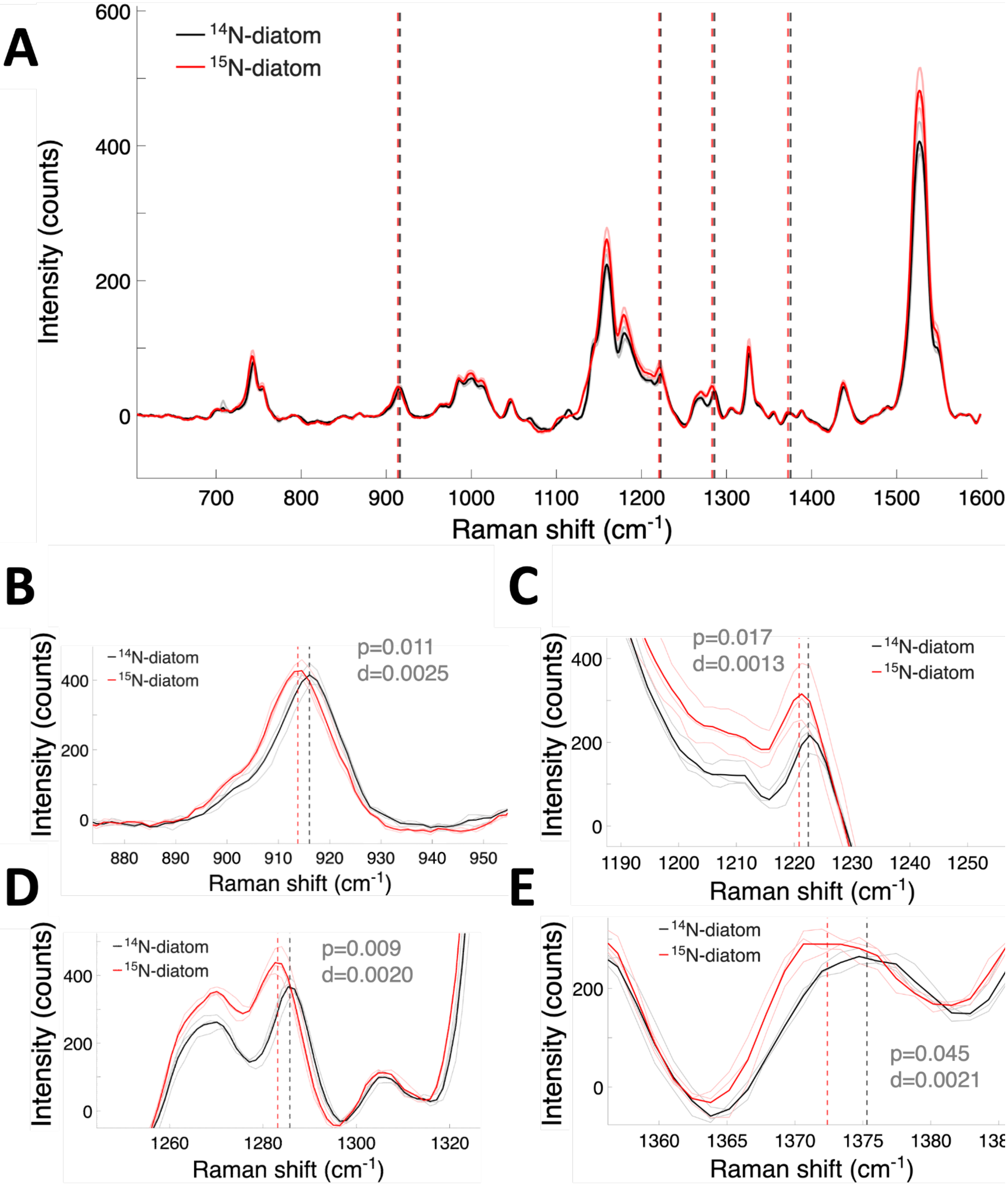


**Figure S3 | Raman spectra of diatoms grown in $^{14}$N and $^{15}$N media reveal four spectral signatures of $^{15}$N incorporation.**

**A:** Mean Raman spectra of C. weissflogii grown in $^{14}$N f/2+Si medium (black) and $^{15}$N f/2+Si medium (red). The 785-nm laser excitation generated data specifically reflecting chlorophyll and carotenoids pigments[45–47]. Thin transparent lines represent the mean Raman spectrum of each of three independent cultures per treatment, with each spectrum obtained by averaging 25 individual diatom cells. Thick lines represent the average spectrum across the three biological replicates for each treatment. Vertical dashed lines indicate the mean positions of the four Raman peaks that exhibited a significant downshift in $^{15}$N-grown diatoms compared with $^{14}$N-grown diatoms (two-sample t-test). **B–E:** Enlarged views of the four shifted peaks. For each peak, the plot reports the p-value (p) from a two-sample t-test and the relative shift in the mean peak position between $^{14}$N- and $^{15}$N-grown diatoms, calculated as $d=(\mu_{14N}-\mu_{15N})/\mu_{14N}$ , where $\mu_{14N}$ and $\mu_{15N}$ are the mean peak positions for $^{14}$N and $^{15}$N-grown diatoms.

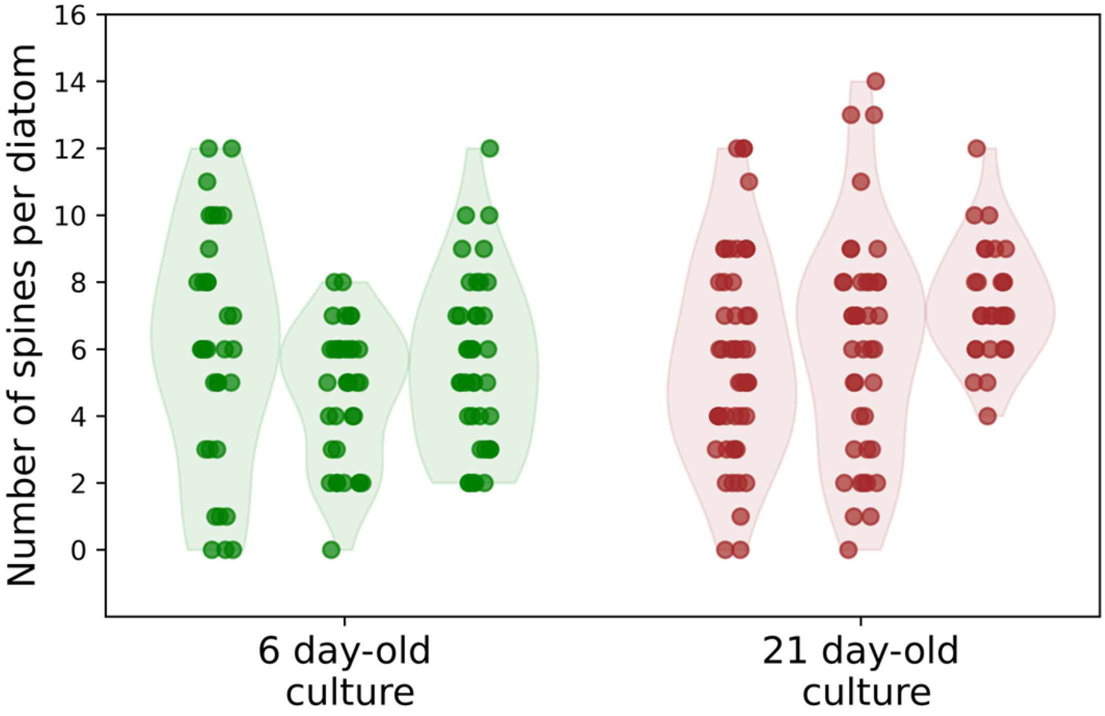


**Figure S4 | The number of spines per diatom is independent of culture growth phase**

The number of spines per cell was quantified in three independent axenic *C. weissflogii* cultures using calcofluor white staining, at 6 days and 21 days post cultures inoculation, showing no significant variation in spine number between growth phases.

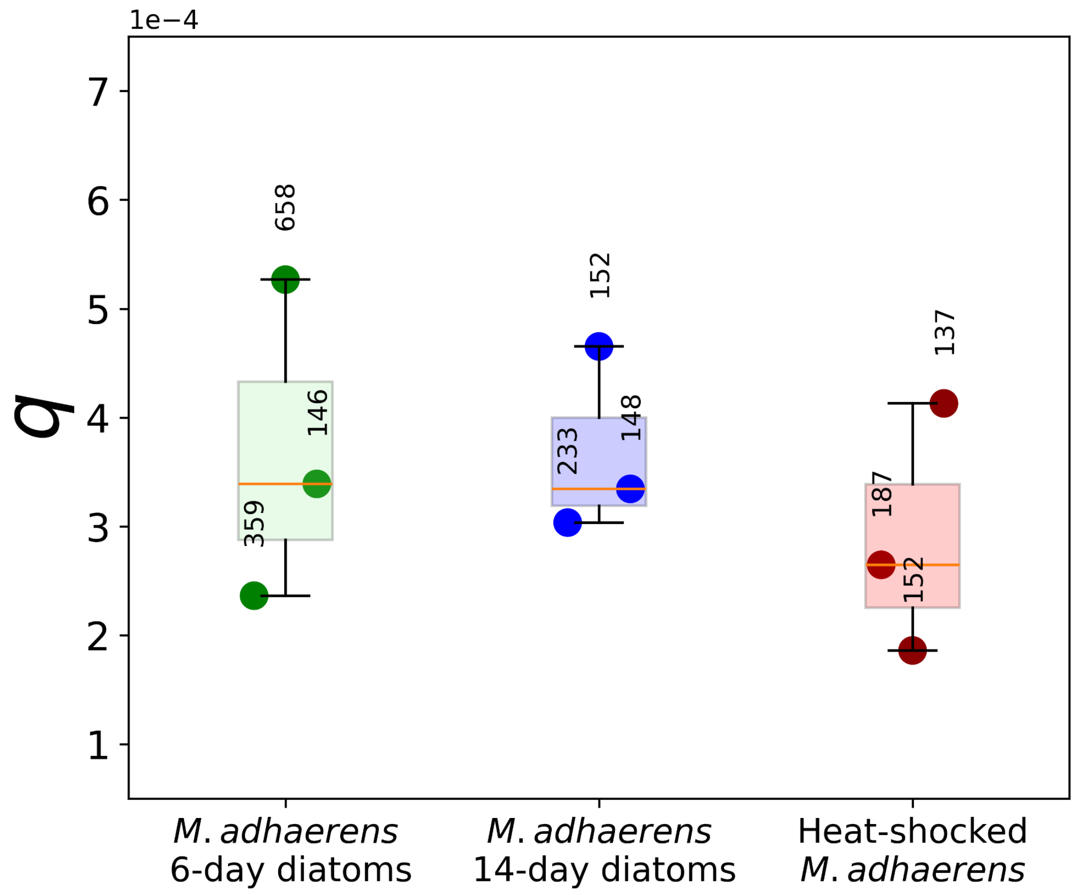


**Figure S5 | Neither diatom growth stage nor bacterial lysis affects bacterial affinity for *C. weissflogii* spines.**

The probability ($q$) of a bacterium attaching to the spines of a diatom upon encounter was measured in triplicate using the microfluidic setup shown in Fig. 4A. In the first condition (green), diatoms were harvested after 6 days of growth; in the second condition (blue), after 14 days of growth. In the third condition (red), bacteria were heat-killed prior to the experiment, and diatoms were harvested after 6 days of growth. Values next above the data points indicate the number of diatoms examined in each experiment. In each boxplot, the central line indicates the median, the box bounds the interquartile range (25th–75th percentiles) and the whiskers extend to the full data range.

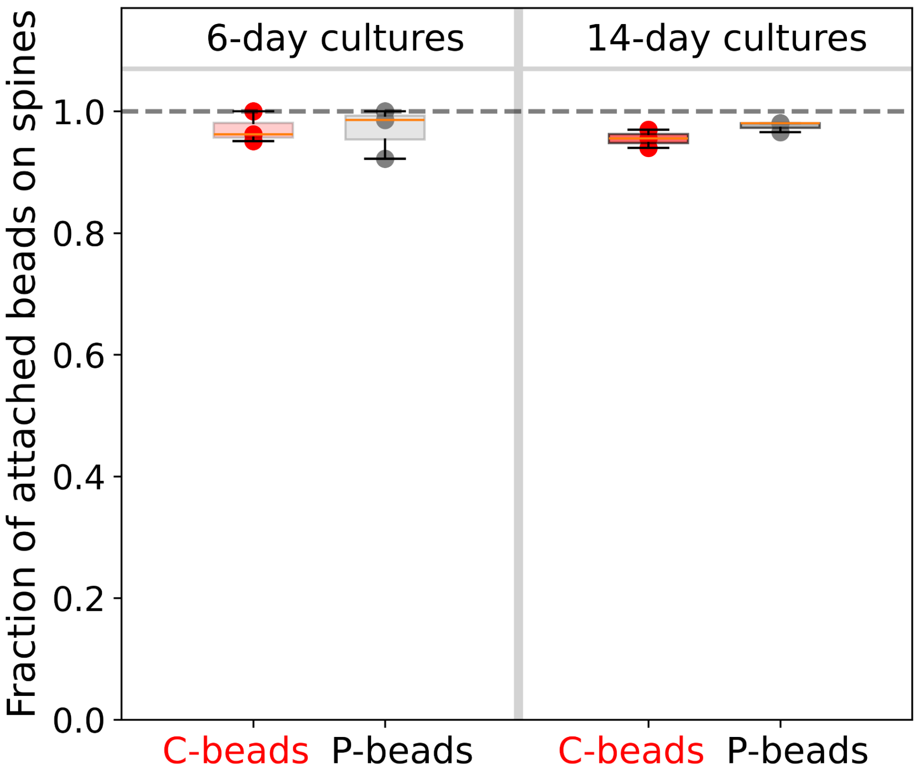


**Figure S6 | Bead attachment in the microfluidic assay occurs predominantly on diatom spines rather than on the cell wall.**

Fraction of bead attachments occurring on diatom spines (as opposed to the cell wall) in the microfluidic experiments shown in Fig. 5. Diatoms from 6-day or 14-day cultures were exposed to carboxylated beads (C-beads) or raw polystyrene beads (P-beads). Experiments were performed in triplicate. Each data point represents the mean fraction of bead attachments occurring on spines in one biological replicate. In each boxplot, the center line indicates the median, the box the interquartile range (25th–75th percentiles), and the whiskers the full data range.

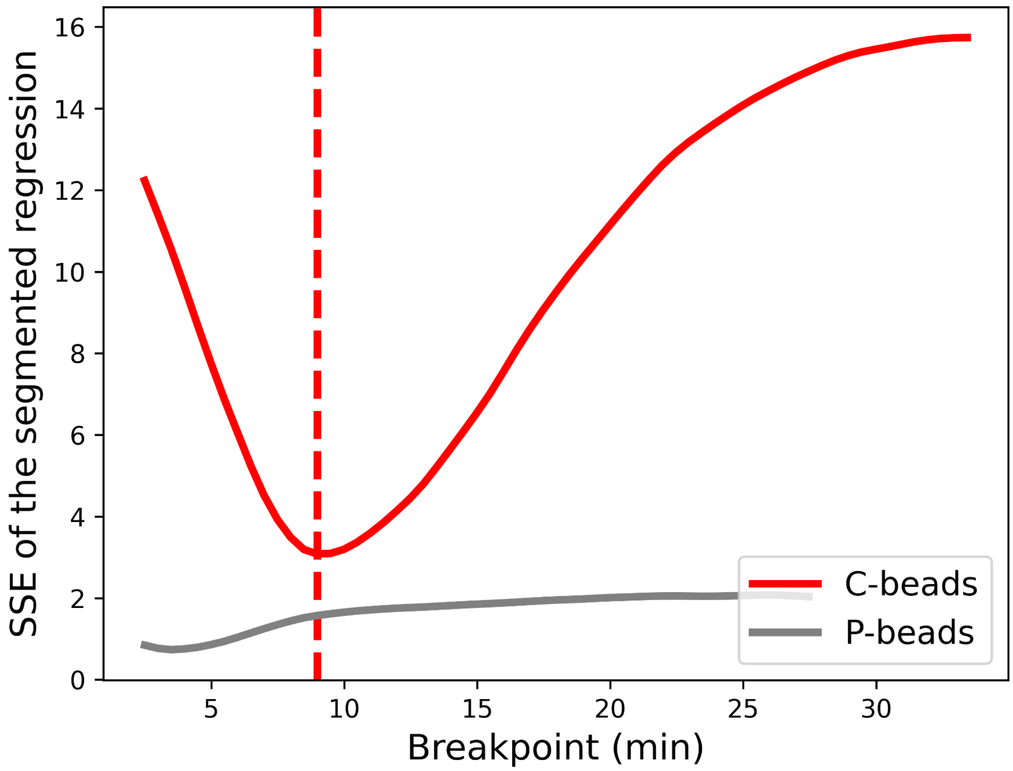


**Figure S7 | Segmented regression reveals a sharp transition in carboxylated bead attachment dynamics to diatom spines after ~9 min of exposure under flow.**

Sum of squared errors (SSE) of segmented regressions fitted to the colonization dynamics (Fig. 3B) as a function of the imposed breakpoint. For each candidate breakpoint, a piecewise linear model was fitted and its SSE computed. The minimum identifies the optimal breakpoint (red dashed line). A pronounced minimum is observed for C-beads (carboxylated polystyrene beads), indicating a clear transition in colonization dynamics, whereas no sharp minimum is detected for P-beads (raw polystyrene beads), consistent with the absence of a distinct regime shift.

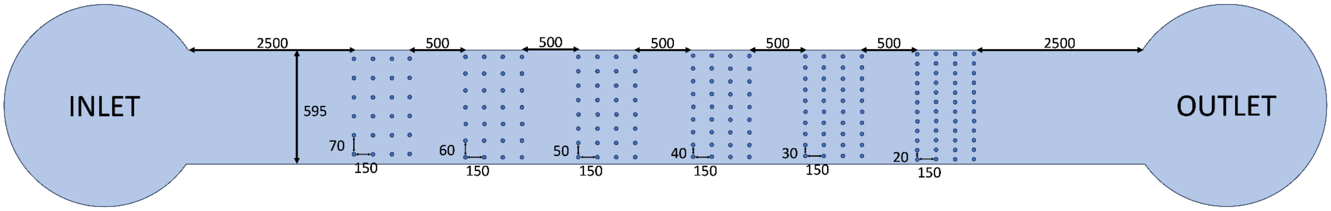


**Figure S8 | Schematic of the microfluidic device.**

All dimensions are in μm. Micropillars have a diameter of 30 μm and the channel height is 106 μm.

**Table S1: Specificity of the primary antibodies used in this study.**

| Primary Antibody | Recognised epitope structure | Reference |
|---|---|---|
| LM11 | (1→4)-β-D-xylan/arabinoxylan | [1] |
| LM21 | (1→4)-β–D-(galacto)(gluco)mannan | [2] |
| JIM13 | Arabinogalactan protein glycan | [3] |
| BAM2 | Sulphated epitope present in sulphated fucan | [4] |

### Note S1: Mechanisms of bacterial lysis on diatom spines

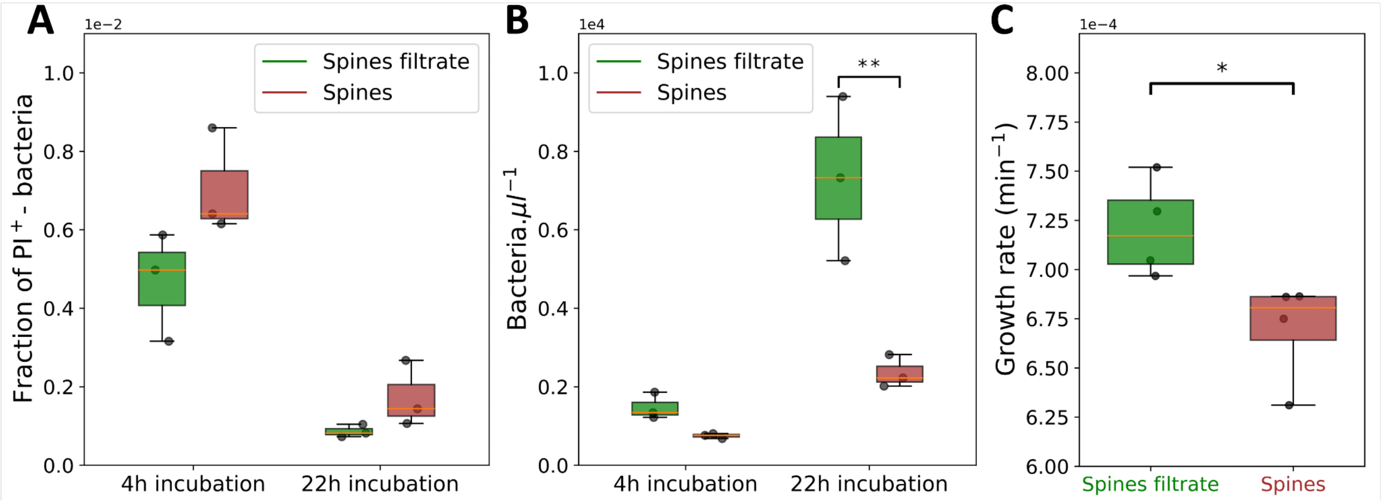


**Figure SN1-1 | Physical interaction with diatom spines imposes physiological stress on *M. adhaerens.***

**A-B:** Fraction of PI-positive (dead) and overall density of *M. adhaerens* cells after resuspension in artificial seawater with either a dense suspension of purified spines (red) or the spine-free filtrate (green). **: Tukey HSD p-value <0.005 **C:** Growth rate of *M. adhaerens* in 10% marine broth supplemented with a dense suspension of purified spines (red) or the corresponding spine-free filtrate (green). *: t-test p-value <0.05. In each boxplot, the horizontal line indicates the median, the box bounds the interquartile range (25th–75th percentiles), and the whiskers extend to the full data range.

The observation that 26.3% (± 6.5%) of spine-attached bacteria from 14-day-old co-cultures stain positive with the membrane-impermeant dye Sytox Green could reflect either a higher propensity of damaged *M. adhaerens* cells to adhere to *C. weissflogii* spines, or bacterial lysis occurring as a consequence of attachment. Two lines of evidence are more consistent with the latter interpretation.
First, SEM and cryo-ET imaging reveal pili-like appendages entwined with diatom spines (Fig. 1, Fig. S1, Fig. S2), suggesting that attachment is at least partially mediated by active pili-driven contact rather than passive adhesion of non-viable cells.
Second, heat-killed *M. adhaerens* cells did not show increased binding affinity to diatom spines relative to healthy cells in the microfluidic assay (Fig. S5), arguing against the possibility that membrane damage per se promotes spine adhesion. Together, these observations are consistent with the interpretation that the observed lysis of 26.3% (± 6.5) of spine-associated bacteria occurs as a

consequence of attachment rather than preceding it, though the temporality and mechanisms of bacterial lysis on diatom spines warrant dedicated future investigation.

Motivated by the detection of fucoidan on the spines (Fig. 6) —a polysaccharide with reported antimicrobial activity[5]— we tested whether the spines themselves exhibit antibacterial properties that could account for the observed lysis of spine-attached bacteria. *M. adhaerens* cells were incubated with a concentrated suspension of isolated spines (~$2 \times 10^7$ spines $mL^{-1}$, equivalent to approximately $2 \times 10^6$ diatom cells; Fig. S4; see Materials and Methods for the isolation protocol) in either artificial seawater (for viability assessment by propidium iodide staining) or marine broth (for growth assays by plate reader). Each experiment included a control in which the spines were removed by 0.2 µm filtration to account for the possible effect of diatom-derived compounds that were not eliminated by the washing steps. We detected a significant effect of spines on the proportion of PI-positive bacteria (two-way ANOVA, $p = 0.029$), with a slightly higher proportion of PI-positive bacteria in the presence of spines compared with the spine filtrate. Pairwise comparisons at individual time points did not reach statistical significance, suggesting limited evidence for an antibacterial effect of diatom spines at the *M. adhaerens* population level. However, a weak population-level signal would be consistent with a mechanism in which spine-mediated killing requires physical attachment. Under such a mechanism, even highly efficient lysis following attachment could produce only a modest population-level increase in PI-positive cells given the rarity of attachment events within the bacterial population. In principle, this hypothesis could be tested by capturing the full sequence of microscale events—from encounter to attachment to lysis—for a large number of bacteria. In practice, however, the infrequency of attachment events makes such an experiment technically very challenging.

In the same experiment, diatom spines significantly reduced bacterial density after 22 h of incubation compared with the spine filtrate (Tukey HSD, $p = 0.002$; Fig. SN1-1B), whereas no significant effect of spines was detected after 4 h ($p = 0.84$). The impact of spines on bacterial population growth was further quantified in marine broth using a plate-reader assay. The presence of spines significantly reduced bacterial growth relative to the spine-filtrate control, demonstrating a bacteriostatic activity (independent two-sample t-test, $p=0.03$, Fig. SN1-1C). To specifically assess the effect of spine-associated fucoidan on *M. adhaerens*, we repeated the viability and growth assays using commercially available fucoidan in place of purified spines. At concentrations comparable to those measured in seawater near fucoidan-producing brown algae (1 $g.L^{-1}$, ref.[6]), fucoidan exerted a bacteriostatic effect on *M. adhaerens* but caused no detectable lysis during the first hours following incubation (Fig. SN1-2).

In summary, although we cannot yet conclusively link bacterial lysis directly to spine attachment, our results demonstrate that interaction with *C. weissflogii* spines imposes a measurable physiological cost on *M. adhaerens*. This cost could –at least partially– be explained by the bacteriostatic action of fucoidan. Such stress may constrain bacterial proliferation on spines—despite the potential advantage of proximity to diatom-derived dissolved organic matter—and promote the release of organic compounds from interacting bacteria, as seen by cryoET. These processes enhance nutrient availability for colonized diatoms, contributing to the ecological benefits of harboring bacteria on their spines.

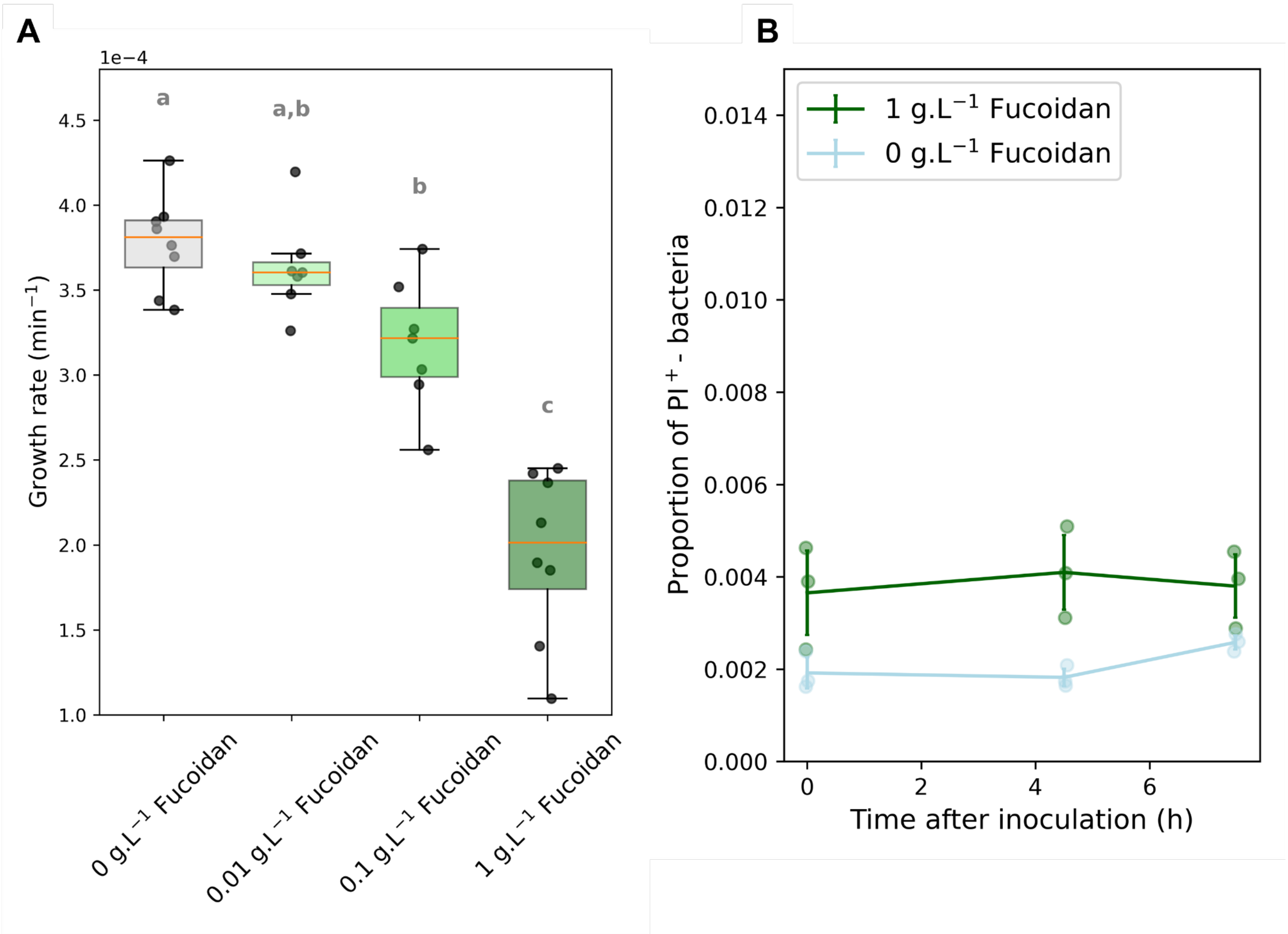


**Figure SN1-2 | Fucoidan exerts no detectable antibacterial activity at the population level but has a bacteriostatic effect on M. adhaerens.**

**A:** Growth rates of *M. adhaerens* in 10% marine broth supplemented with varying fucoidan concentrations. Different letters above boxplots indicate statistically significant differences among treatments, as determined by Tukey's HSD post-hoc test following one-way ANOVA. In each boxplot, the orange line indicates the median, the box bounds the interquartile range (25th–75th percentiles), and the whiskers extend to the full data range.
**B:** Fraction of PI-positive *M. adhaerens* cells over time with 1 g.L$^{-1}$ fucoidan.

### Note S2: Encounter kernel between a bacterium and the spines of a diatom under various ecological conditions

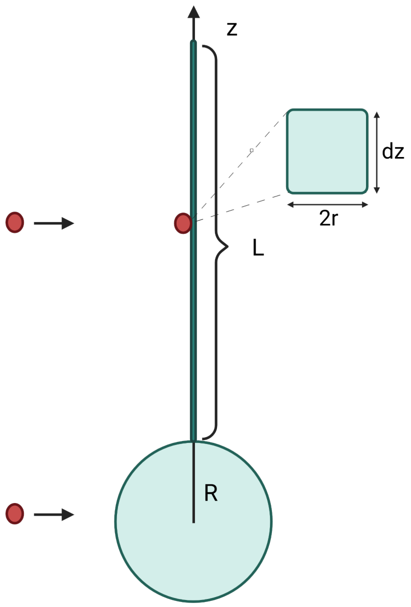


**Figure SN2 | Spines are modelled as infinitely thin lines and bacteria as spheres of radius r.** The collision cross section between a short spine section of length dz and a bacterium is thus 2rdz (see inset).

To get an order-of-magnitude estimate of the collision rate between the diatom spines and bacteria for different encounter mechanisms, we modeled a diatom spine as a thin line and a bacterium as a sphere of radius *r* (Fig. SN2). The encounter kernel of a bacterium and a spine can then be calculated by integrating the encounter kernels along the length of the spine, from *R* (the diatom radius) to *R*+*L*, where *L* is the average spine length. Building on standard encounter rate theory[7–9], which describe encounter kernels between objects under diffusion-, flow-, motility-, and gravitational sinking–driven regimes, we derived simplified expressions for encounters between a bacterium and a diatom spine. The expressions neglect hydrodynamic interactions between bacteria and the diatom cell body and assume that the cell body and spines intercept bacteria independently from each other. We stress that the calculations below are simplifications aimed at estimating the order-of-magnitude of the collision rates.

*Encounters driven by turbulent flow*

The encounter kernel between non-motile bacteria and the diatom cell body under turbulent flow is given by: $\Gamma_{flow_{cb}} = 1.3(\epsilon/\nu)^{1/2}(r+R)^3$, where $\epsilon$ is the kinetic energy dissipation rate, $\nu$ the kinematic viscosity of seawater, $r$ the bacterial radius, and $R$ the diatom cell radius.

To describe the encounter kernel between non-motile bacteria and a single diatom spine, we account for the local flow velocity gradient along the spine. We approximate the turbulent flow by a simple shear flow, where the flow velocity v(z) at a distance z from the diatom surface scales as $v(z) = z(\epsilon/\nu)^{1/2}$.

Assuming the spine is oriented in the direction perpendicular to the flow, we have: $\Gamma_{flow_{sp}} = 2r\int_R^{L+R} v(z)dz = 2r(\epsilon/\nu)^{1/2}\int_R^{L+R} zdz = r(\epsilon/\nu)^{1/2}((R+L)^2 - R^2)$

*<u>Encounters driven by bacterial motility</u>*

In the case of swimming bacteria, the encounter kernel describing encounters driven by bacterial motility and a stationary diatom cell body can be expressed as: $\Gamma_{mot_{cb}} = \pi(r+R)^2 f_{mot}U$, where $U$ is the average bacterial swimming speed and $f_{mot}$ is the fraction of motile cells within the bacterial population. Similarly, the encounter kernel between bacteria and a stationary diatom spine can be obtained by averaging over random bacterial approach angles. This yields: $\Gamma_{mot_{sp}} = 0.25\pi\int_R^{L+R} f_{mot}Udz2r = 0.5\pi r f_{mot}U\int_R^{L+R} dz = 0.5\pi r f_{mot}UL$

The prefactor 0.25 accounts for the fact that bacterial trajectories are not uniformly perpendicular to the spine surface, which reduces the effective encounter cross-section. For marine bacteria, we use $f_{mot}$~0.5 [10,11] and $U$~50 µm s$^{-1}$ (ref. [11]).

*<u>Encounters driven by diatom sinking</u>*

A stationary, sinking diatom cell body generates a Stokes flow in its vicinity, which influences its encounter rate with surrounding bacteria. Under these conditions, the encounter kernel between bacteria and sinking stationary diatom cell body with sinking speed $V$ is given by $\Gamma_{sink_{cb}} = 1.45\pi V r^2$ (ref. [7]07/09/2026 12:44:00).

Similarly, the encounter kernel between bacteria and a stationary diatom spine can be obtained by averaging over random bacterial approach angles and neglecting hydrodynamic interactions between the spine and the bacteria, which yields: $\Gamma_{sink_{sp}} = 0.25\pi\int_R^{L+R} V2rdz = 0.5\pi rV\int_R^{L+R} dz = 0.5\pi rVL$

Diatom spines reduce the drag of the cell[12–14], thereby affecting its sinking speed $V$. We thus computed $V$ as a function of $L$ and $n$, the number of spines per diatom cell. We consider a perfectly spherical diatom of radius *R* sinking at very low Reynolds number. The sphere carries *n* rigid, straight spines, each of length *L* and diameter *d*, with *d* much smaller than *L*. Hydrodynamic interactions between spines are neglected.

According to Stokes' law, the drag force ($F_D$) acting on a sinking cell moving at velocity $V$ is $F_D = 6\pi\mu RV$ , where $\mu$ is the dynamic viscosity of sea water[15]. At terminal velocity $V_0$, the drag force balances the net gravitational force $F_g$. Therefore, the terminal sinking velocity of a spineless phytoplankton cell is $V_0 = \frac{F_g}{6\pi\mu R}$.

For a spine, the drag coefficient parallel to its axis can be expressed as $\zeta_{parallel}(L) = \frac{2\pi\mu L}{ln(L/d)-0.5}$ and the drag coefficient perpendicular to its axis as $\zeta_{perpendicular}(L) = \frac{4\pi\mu L}{ln(L/d)+0.5}$ (ref.[16]).

For an arbitrary angle $\theta$ between the spine axis and the flow, the instantaneous drag coefficient of a spine $\zeta_{spine}(L,\theta)$ can be approximated by decomposing the flow velocity into components parallel and perpendicular to the spine. This leads to a linear decomposition of the drag coefficient as $\zeta_{spine}(L,\theta) = cos^2(\theta)\zeta_{parallel}(L) + sin^2(\theta)\zeta_{perpendicular}(L)$ , where the squared trigonometric terms represent the projection of the flow velocity onto the spine axis and its normal directions. Assuming a uniform distribution of spines orientations, the averages $< cos^2(\theta) >= 1/3$ and $< sin^2(\theta) >= 2/3$ yield the average drag coefficient of a spine $\zeta_{spine}(L) = \frac{1}{3}\zeta_{parallel}(L) + \frac{2}{3}\zeta_{perpendicular}(L)$ .

Under the assumption of no hydrodynamic interaction between the diatom cell body and its spines, the sinking velocity *V(n,L)* of a diatom harboring *n* spines can be expressed as $V(n,L) = \frac{F_g}{\zeta_{total}(n,L)}$ where:

$\zeta_{total}(n,L) = 6\pi\mu R + n\zeta_{spine}(L)$

This gives: $\frac{V(n,L)}{V_0} = \frac{1}{1+n\zeta_{spine}(L)/6\pi\mu R}$, which can be explicitly written as:

$$V(n,L) = \frac{V_0 6R}{6R+nL(\frac{2}{3(ln(L/d)-0.5)}+\frac{8}{3(ln(L/d)+0.5)})}.$$

We use $V_0 = 1\ \mu m.s^{-1}$ as the representative sinking velocity of a spineless *C. weissflogii* cell based on experimental measurement across multiple *Thalassiosira* species[17].

<u>*Encounters driven by bacterial diffusion:*</u>

Finally, the encounter kernel driven by bacterial diffusion between bacteria and a stationary diatom cell body can be expressed as: $\Gamma_{diff_{cb}} = 4\pi(r + R)D_b$, where $D_b$ is the diffusion coefficient of bacteria.

Following the formulation of Berg[8] for diffusive encounters with an elongated ellipsoid verifying $(\frac{L}{2})^2 >> d^2$, the diffusion-driven encounter kernel between bacteria and a diatom spine of length *L* and diameter *d* can be approximated as: $\Gamma_{diff_{sp}} = 2\pi * D_b L / (ln(\frac{L}{d}))$ .

We estimated the encounter rate (*E)* between bacteria and the spines of a diatom or its cell body for each encounter mechanism as $E = C_b \Gamma$, with $C_b$=$10^6$ bacteria $mL^{-1}$ (Fig. 5).

### Note S3: Calculation of attachment probability based on experimental data.

We aim at measuring $q$, the probability that a bacterium attaches to the spines of a diatom upon an encounter.

$q$ is given by: $A = \mathrm{q} C_d C_b \Gamma_{sp}$ *Equation N3.1*

where $A$ is the rate of attachment events to the diatom spines per unit of volume, $C_d$ *and* $C_b$ are the concentration of diatom and bacteria, respectively, and $\Gamma_{sp}$ the kernel of bacteria-diatom spines encounter.

In our experiments, we determined $A$ by quantifying the temporal change in the concentration of bacteria attached to diatom spines. Specifically, we defined:

$A = dC_{a\,b}/dt \; = \; [C_{ab}\,(t_{exp}) - C_{ab}(t_0)]/t_{exp}$ *Equation N3.2*

where $C_{ab}$ is the concentration of attached bacteria and $t_{exp}$ is the duration of the experiment.

This framework was applied to both the microfluidic attachment assays and the coculture experiments to estimate $q$ under each experimental context.

*Microfluidic experiment:*

In the microfluidic chip, we typically observed no more than one bacterium attached per diatom spine by the end of the experiment. The concentration of attached bacteria can therefore be approximated by the concentration of colonized diatoms. The concentration of colonized diatoms $C_{cd}$ is obtained from the experimentally measured fraction of colonized cells, $f_{col}$, such that: $C_{cd}(t) = f_{col}(t) C_d$

Substituting this into Equation N3.2 yields:

$A = [f_{col}(t_{exp}) - f_{col}(\mathrm{t} = 0)]\; C_d/t_{exp}$ *Equation N3.3*

and combining with Equation N3.1 gives:

$q = [f_{col}(t_{exp}) - f_{col}(\mathrm{t} = 0)]/(t_{exp} C_b \Gamma_{sp})$ *Equation N3.4*

In the microfluidic chip, bacterial encounters with the spines of diatoms immobilized between micropillars are generated by pump-driven flow. The flow rate was set to match the swimming speed of *M. adhaerens*[18] (~20 µm s⁻¹), and effective flow within the channel was verified by tracking fluorescent beads. Measured flow velocities fluctuated by a factor of 2–3 around the expected value but averaged ~20 µm s⁻¹, consistent with the target speed (Fig. SN3-1). Under these conditions, encounters between bacteria and diatom spines in the chip can be regarded as equivalent to a diatom sinking at 20 µm s⁻¹ through a water column with bacterial concentration $C_b$.

As detailed in Note S2, the encounter kernel $\Gamma_{sp}$ between a bacterium (with radius $r$=1 µm) and the spine of diatom with length $L$=100 µm sinking at a rate of $V$ can be expressed as: $\Gamma_{sp} = \; 0.5\pi r V L$.

From the bacterial perspective, the spines of a diatom can be treated as independent encounter sites. The corresponding encounter kernel between a bacterium and the spines of a diatom is therefore given by $\Gamma_{sp} = 0.5\pi\text{nrVL}$, where *n* is the number of spines per diatom cell. *n* was set to *n* = 10 based on experimental measurements (Fig. S4).

Equation N2.4 therefore becomes

$$q = \left[f_{col}\left(t_{exp}\right) - f_{col}(\text{t} = 0)\right] / (t_{exp} C_b 0.5\pi nrVL) \qquad \text{Equation N3.5}$$

In the microfluidic chip, the bacterial concentration was constant at 5x10$^6$ cells mL$^{-1}$, and the experiment duration was $t_{exp}$= 30 min. Because no bacteria were present in the chip prior to flow initiation, $f_{col}(\text{t} = 0) = 0$.

For *M. adhaerens,* Equation N3.5 yields *q* ~ 3.7x10$^{-4}$.

Applying the same calculation to *Alteromonas macleodii* and several *Vibrio* species (Fig. 4B) produced similar estimates of *q*, indicating that attachment probability per encounter is comparable across diverse bacterial taxa interacting with *C. weissflogii*.

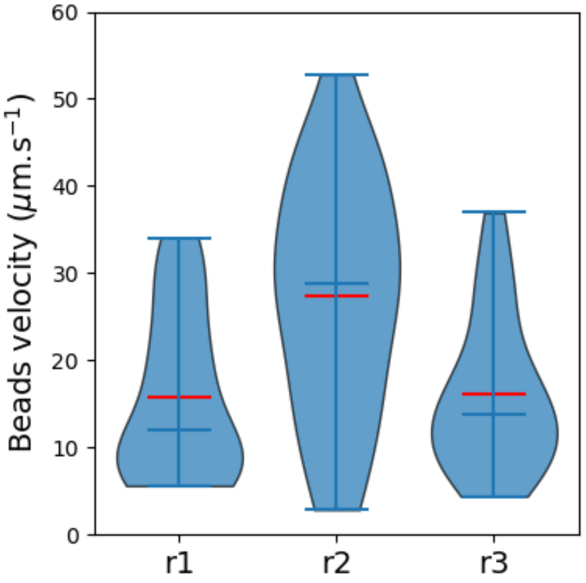


**Figure SN3-1 | Distribution of beads velocity in the microfluidic chip.** The experiment was performed in triplicates. The trajectory of approximately 30 beads was recorded in each experiment. The red lines indicate the mean bead velocity while the horizontal blue lines indicate the median bead velocity.

<u>*Coculture experiments:*</u>

At days 6 and 13 after coculture inoculation, bacterial and diatom abundances were quantified by flow cytometry (Fig. SN3-2). The number of bacteria found attached to diatom spines was measured across hundreds of cells by epifluorescence microscopy on the >8 µm size fraction.

The rate of attachment to diatom spines, *A*, was calculated following Equation N3.2:

$$A = [C_{ab}(t_{stat}) - C_{ab}(t_{exp})] / (t_{stat} - t_{exp})$$

In these experiments, we quantified the mean number of attached bacteria per diatom, $N_{bpds}$, and therefore express the attachment rate as:

$$A = C_d * [N_{bpds}(t_{stat}) - N_{bpds}(t_{exp})]/(t_{stat} - t_{exp}) \qquad \textit{Equation N3.6}$$

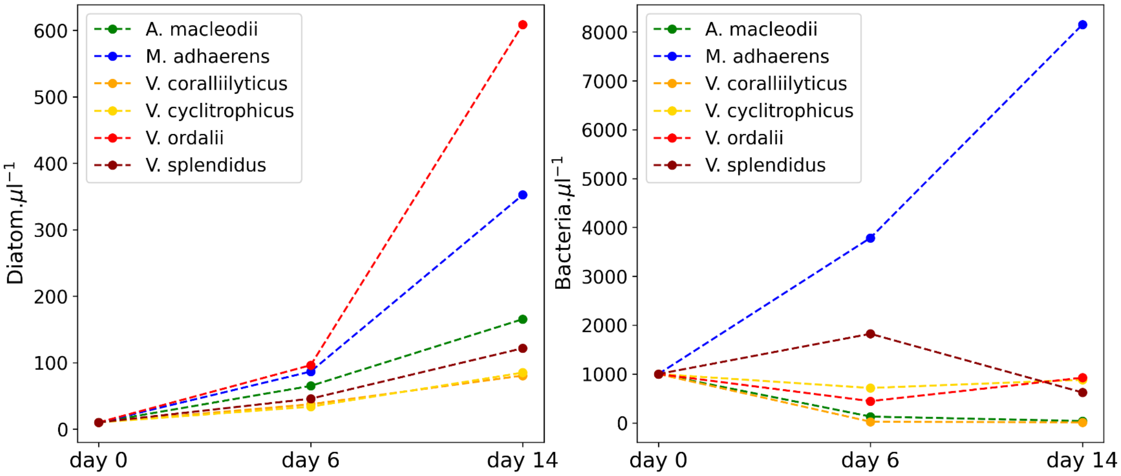


**Figure SN3-2 | *C. weissflogii* and bacterial densities at day 6 and day 11 following coculture inoculation.**

Because cocultures were grown without shaking, we considered bacterial motility to be the dominant driver of encounters between bacteria and diatom spines. As detailed in Supplementary Note S2, the corresponding encounter kernel is: $\Gamma_{mot_{sp}} = 0.5\pi r f_{mot} U L n$, where *n* is the number of spines per diatom, $f_{mot}$ the fraction of motile bacteria and *U* the bacteria speed.

Combining Equation N3.1 with Equation N3.6 yields:

$$q = [N_{bpds}(t_{stat}) - N_{bpds}(t_{exp})]/((t_{stat} - t_{exp}) C_b 0.5\pi r f_{mot} U L n)$$

We applied this calculation to data from a range of bacterial species (Fig. SN3-3). All tested bacterial strains exhibited low *q* values, consistent with the microfluidics experiment. For *M. adhaerens*, we obtained $q \sim 7 \times 10^{-6}$.

The 50-fold difference in *q* values obtained in the two experimental set-ups may result from processes that manifest only in the coculture experiment over longer timescales (>30 min) such as bacterial detachment from phytoplankton or modulation of bacterial attachment by accumulated diatom exudates.

Finally, we estimated the number of encounters $N_e$ between bacteria and the spines of a single diatom over a duration $T$=1h at day 6 post-inoculation. This quantity can be expressed as: $N_e = \Gamma_{mot_{sp}} C_b T$ . Using the mean bacterial concentration measured at day 6 across three independent cocultures, we obtained $N_e$ ~ 380 bacterial encounters per diatom per hour.

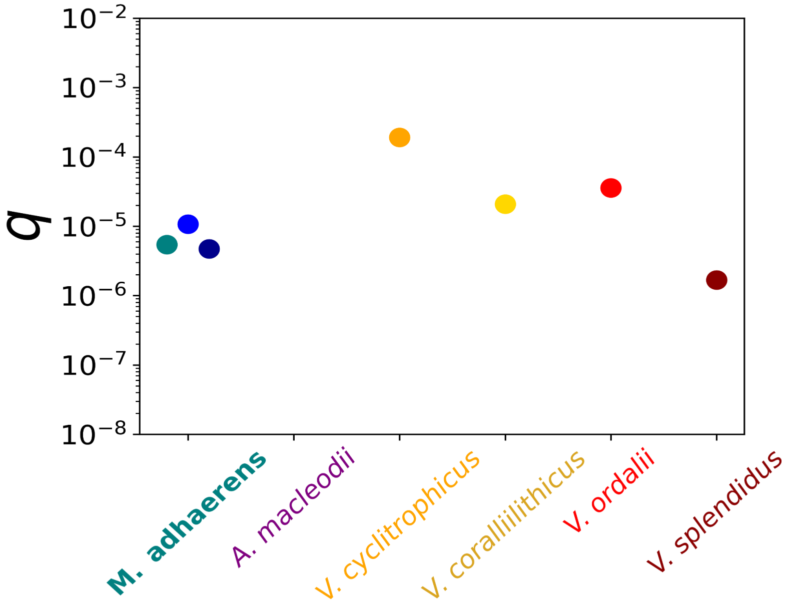


**Figure SN3-3 | Attachment probability q of various bacterial species to *C. weissflogii* spines upon an encounter estimated in cocultures.** No data point is shown for *A. macleodii* because the calculated q value was negative. As q represents a probability, negative values are not physically meaningful and arose here from a higher number of bacteria per diatom spine at 6 days than at 14 days of coculture (although the values were very low in both cases).

### Note S4: Prediction of bacterial attachment rate to diatom spines

To estimate the rate of bacterial attachment to diatom spines under bloom conditions, we considered encounters driven independently by turbulent flow, bacterial motility, diatom sinking and diffusion. We approximate the total encounter kernel $\Gamma_{sp}$ as the sum of the encounter kernels of these four mechanisms[19], thus:

$$\Gamma_{sp} = n(r(\epsilon/\nu)^{1/2}R^2((1+L/R)^2-1) + 0.5\pi r f_{mot}UL + 0.5\pi r V(n,L)L + 2\pi D_b L/(ln(L/d)))$$

*Equation N4.1*

where $n$ is the number of spines per diatom cell, $r$ is the bacterial radius, $R$ the diatom radius, $L$ the spine length, $d$ the spine radius, $U$ the bacterial swimming speed, $f_{mot}$ the fraction of motile bacteria, $V$ the diatom sinking velocity, $\varepsilon$ the turbulent kinetic energy dissipation rate, $\nu$ the kinematic viscosity of seawater and $D_{bact}$ the diffusion coefficient of bacteria.

The attachment rate $A$, as defined in Equation N3.1, is then given by:

$$A = qn[(r(\epsilon/\nu)^{1/2}R^2((1+L/R)^2-1) + 0.5\pi rL(f_{mot}U + V(n,L)) + 2\pi D_b L/(ln(L/d))]C_b C_d$$

*Equation N4.2*

where $q$=3.7x$10^{-4}$ represents the experimentally determined probability that a bacterium attaches upon encountering the spines of a diatom cell.

Assuming $d = 100\,nm, n = 10, f_{mot} = 0.5, D_b = 10^{-13}\,m^2.s^{-1}, U = 50\,\mu m.s^{-1}, V_0 = 1\,\mu m.s^{-1}$ (ref.[17]), $\nu = 10^{-6}\,m.s^{-2}$ and microbial abundance of $C_b$ and $C_d$ constant and equal to $10^6$ bacteria $mL^{-1}$ and $10^5$ diatoms $mL^{-1}$ over the stationary phase of a bloom[20–23], we computed the attachment rate $A$ across a range of spine lengths and turbulence intensities (Fig. SN4). Spine lengths were varied from values comparable to a single cell body ($L$ = 1–10 µm) to approximately 30 cell bodies ($L$ = 300 µm). The range of turbulent kinetic energy dissipation rates ($\varepsilon$ = $10^{-8}$–$10^{-4}$ W $kg^{-1}$) was based on field measurements[24,25]. Diatoms are expected to experience high $\varepsilon$ values near the surface due to wave-driven mixing, and progressively lower $\varepsilon$ values as they sink into deeper layers of the water column.

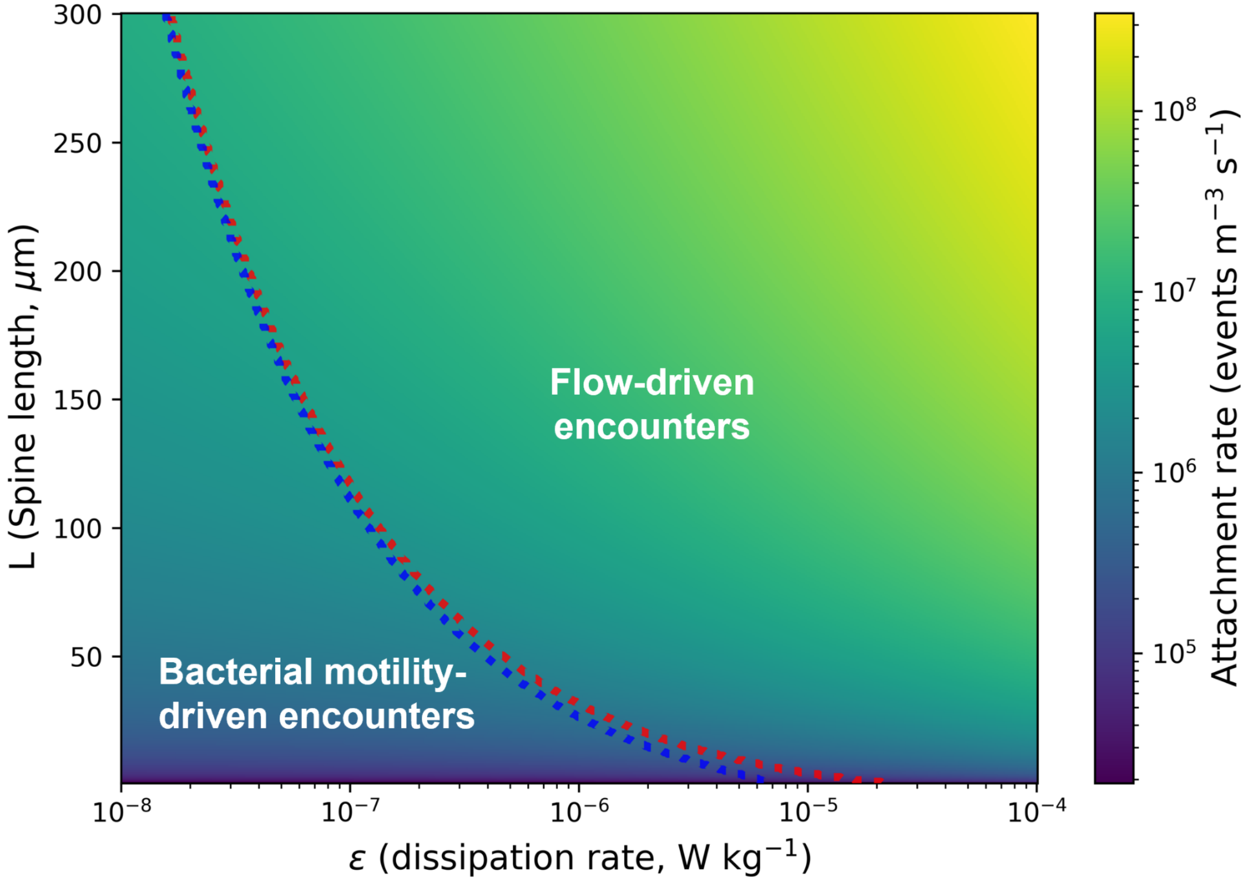


**Figure SN4 | Modeled bacterial attachment rate to diatom spines as a function of turbulent kinetic energy dissipation rate (*ε*) and spine length (*L*).**
Color shading denotes the attachment rate (events $m^{-3}$ $s^{-1}$). The red dotted contour marks the transition between two encounter regimes: above the contour (higher ε for a given L), encounters are dominated by turbulence ($\Gamma_{flow_{sp}} > \Gamma_{mot_{sp}} + \Gamma_{sink_{sp}} + \Gamma_{diff_{sp}}$). The blue dotted contour indicates a second regime transition: below this contour, encounters are driven primarily by bacterial motility ($\Gamma_{mot_{sp}} > \Gamma_{flow_{sp}} + \Gamma_{sink_{sp}} + \Gamma_{diff_{sp}}$). Model parameters are n=10, R = 5 μm, r = 1 μm, d=100 nm, $D_{bact}$ = $10^{-13}$ $m^2$ $s^{-1}$, U = 50 μm $s^{-1}$, $f_{mot}$ =0.5 and $V_0$ = 1 μm $s^{-1}$.

This diagram illustrates how hydrodynamic and morphological factors jointly control colonization potential during the stationary phase of a bloom. Bacterial attachment rates are highest for diatoms bearing long spines in surface waters, where turbulence levels are high ($\varepsilon$ = $10^{-5}$-$10^{-4}$ W $kg^{-1}$ ref.[24,25]). Under these conditions, strong turbulent shear enhances bacterial interception by diatom spines, which extend away from the cell body into regions of higher flow velocity. For $L$ = 100 μm and $\varepsilon = 10^{-4}$ W kg$^{-1}$, the model predicts an attachment rate corresponding to $36\times10^{11}$ attachment events per cubic meter after one day of stationary bloom. Assuming these attachment events are uniformly distributed across the diatom population, each diatom cell would host on average 36 bacteria on its spines after one day of stationary bloom.

As diatoms sink into deeper layers of the water column, characterized by lower $\varepsilon$ values ($\varepsilon = 10^{-8}$-$10^{-7}$ W kg$^{-1}$), they transition into a regime in which bacterial motility becomes the dominant driver of encounters between bacteria and diatom spines. Overall attachment rates are approximately tenfold lower (for $L$ = 100 μm and $\varepsilon = 10^{-7}$ W kg$^{-1}$) in these environments than those predicted for surface waters. After one day of a stationary bloom, the model predicts an average of 2.4 spine-attached bacteria per diatom.

Across the explored parameter space, encounter kernels associated with diatom sinking or bacterial diffusion never exceed those arising from the other encounter mechanisms. Although spines are effective at intercepting small particles during sinking because of their small radius relative to bacterial size, their overall contribution to encounters is limited. The increased drag imposed by spines reduces the sinking velocity of the diatom, thereby decreasing the volume of water sampled by the sinking cell. For diffusion-driven encounters, the extreme slenderness of the spines constrains their effective capture radius, rendering this mechanism negligible relative to encounters driven by turbulent flow or bacterial motility.

### Note S5. Stoichiometric estimation of diatom nitrogen uptake from bacteria attached to their spines.

To evaluate the potential contribution of attached bacteria to the nitrogen requirements of *Conticribra weissflogii*, we first estimated the nitrogen mass of a *C. weissflogii* cell, assuming a spherical geometry with a diameter of 10 µm, corresponding to a cell volume of 524 µm³. Using the biovolume–carbon relationship established for diatoms by Menden-Deuer and Lessard[26], the carbon content is estimated at 46 pg C per cell. Applying the canonical Redfield C:N ratio of 106:16 (ref.[27]) yields a nitrogen content of approximately 8.1 pg N per diatom cell.

The nitrogen content of *Marinobacter adhaerens* was approximated as 8.1 fg N per cell, based on the average measurement of carbon content and C:N ratio of coastal and oceanic marine bacteria reported by Fukuda *et al.*[28]

Assuming a diatom division time of 12 hours[29], the biomass of a diatom population doubles over this period. Each diatom cell must therefore acquire nitrogen equivalent to its initial nitrogen content to support this growth, corresponding to a nitrogen requirement of about 8.1 pg N every 12 hours, or roughly 0.7 pg N $h^{-1}$.

If all nitrogen from a bacterium attached to a diatom spine were fully assimilated by the diatom host, a single diatom would need to capture 88 bacteria per hour to meet its nitrogen demand. This number would be substantially higher if nitrogen assimilation from attached bacteria were incomplete.
